# Unique Amphipathic *α*-helix Drives Membrane Insertion and Enzymatic Activity of ATG3

**DOI:** 10.1101/2023.02.11.528101

**Authors:** Taki Nishimura, Gianmarco Lazzeri, Noboru Mizushima, Roberto Covino, Sharon A. Tooze

**Author notes:** Correspondence: Taki Nishimura, Roberto Covino. These authors contributed equally (Taki Nishimura, Gianmarco Lazzeri).

## Abstract

Autophagosome biogenesis requires a localized perturbation of lipid membrane dynamics and a unique protein-lipid conjugate. Autophagy-related (ATG) proteins catalyze this biogenesis on cellular membranes, but the underlying molecular mechanism remains unclear. Focusing on the final step of the protein-lipid conjugation reaction, ATG8/LC3 lipidation, we show how membrane association of the conjugation machinery is organized and fine-tuned at the atomistic level. Amphipathic *α*-helices in ATG3 proteins (AH_ATG3_) are found to have low hydrophobicity and to be less bulky. Molecular dynamics simulations reveal that AH_ATG3_ regulates the dynamics and accessibility of the thioester bond of the ATG3∼LC3 conjugate to lipids, allowing covalent lipidation of LC3. Live cell imaging shows that the transient membrane association of ATG3 with autophagic membranes is governed by the less bulky- hydrophobic feature of AH_ATG3_. Collectively, the unique properties of AH_ATG3_ facilitate protein-lipid bilayer association leading to the remodeling of the lipid bilayer required for the formation of autophagosomes.

**Teaser:** We uncover the unique biophysical property of amphipathic *α*-helix essential for autophagy

## Introduction

Autophagy is a fundamental cellular event that maintains cellular homeostasis by degrading cytoplasmic materials and damaged organelles. Upon autophagy induction, a cup-shaped membrane structure sequesters a part of cytoplasm and/or selective cargoes by closing to form a double-membrane autophagosome (*1*). Autophagosome formation is driven by the autophagy-related (ATG) proteins and accompanied by dynamic membrane remodeling processes, such as membrane nucleation, expansion and shaping (*2–6*). Several functional groups composed of core ATG proteins play a central role in autophagosome biogenesis: the ULK protein kinase complex, ATG9-containing vesicles, autophagy-specific PI3K (PI3KC3-C1) complex, ATG2-WIPI complex, ATG12–5-16L1 complex, ATG3 and ATG8 proteins (*2, 7*). At the initiation step, multiple ULK complexes are assembled in close apposition to the ER membrane and recruit ATG9 vesicles that act to initiate the seed membrane of the autophagosome precursor. After phosphatidylinositol 3-phosphate (PI3P) generation by PI3KC3-C1 complex, the seed membrane is physically linked to the ER membranes via ATG2-WIPI complex and becomes the destination of ATG2-dependent bulk transport of lipids. Subsequently, membrane lipid imbalance between outer and inner leaflets caused by transferred lipids is equilibrated by the lipid scramblase activity of ATG9 in the nascent autophagosome. This cooperative lipid transport by ATG2 and ATG9, facilitated by a direct interaction between ATG2 and ATG9 (*8*), is thought to induce membrane growth of autophagosomes (*9, 10*). In parallel, ATG8 proteins (LC3s and GABARAPs in mammals) contribute to membrane expansion by different mechanisms. ATG8s undergo lipidation by ATG3 in coordination with the ATG12–5-16L1 complex allowing them to be anchored into autophagic membranes to promote membrane expansion, possibly by inducing membrane hemifusion (*11*) and membrane deformation (*12*). In a recent study, a theoretical model predicts that curvature generation by membrane-shaping proteins including ATG proteins can be a key factor for autophagosome size regulation (*13, 14*).

To accomplish this complicated membrane-mediated process and successfully form autophagosomes, ATG proteins need to interact with membranes in a strictly controlled manner and sense local membrane environments during membrane reorganization. Indeed, ATG proteins have defined spatiotemporal recruitment patterns and specific distributions on autophagic membranes (*15, 16*). This is presumably because each functional group of ATG proteins associates with membranes by a distinct mechanism (*7*). Among lipid-binding modules, amphipathic *α*-helices (AHs), which sense membrane charge, curvature, unsaturation and lipid composition (*17*), are commonly found in some ATG proteins: ATG14L, VPS34, ATG2, WIPIs, ATG3 and ATG16L1 (*18–22*). Interestingly, PI3KC3-C1 containing ATG14L and VPS34 localizes to both the ER membranes and autophagic membranes (*23, 24*), while ATG2 and ATG12–5-16L1 are detected at the edge of growing phagophores (*13, 25–27*), suggesting that individual AH_ATG_ sense a different membrane environment and have a distinct role in autophagosome formation.

Upon nutrient starvation or autophagy stimulation, ATG8/LC3 proteins undergo lipidation by the E1-E2-E3-like enzymatic cascade. The E2-like ATG3 directly binds to membranes via its N-terminal AH_ATG3_ to transfer ATG8/LC3 to phosphatidylethanolamine (PE) (*22, 28, 29*) by bimolecular nucleophilic substitution SN2 reaction (**Fig. S1A**) (*30, 31*). As the AH_ATG3_ shows a preference for highly curved membranes *in vitro* (*32*) and AH_ATG3_ is indispensable for LC3 lipidation (*22*), ATG3 could execute LC3 lipidation on highly curved autophagic membranes. Despite such findings, how ATG3 recognizes membrane lipids via its AH_ATG3_ and the dynamics of ATG3 on autophagic membranes are poorly understood. Whether membrane binding and curvature sensing are the exclusive functions of AH_ATG3_ in autophagy is still unresolved. In fact, ATG16L1, a component of the E3-like ATG12–5-16L1 complex, is a dominant factor for curvature sensing among the components of LC3 conjugation machinery (*19, 33*), implying that AH_ATG3_ has another pivotal function in addition to membrane anchoring (*34*).

To address the function of ATG3 in autophagy we focused our studies on the essential lipid binding module AH_ATG3_. Unsupervised machine learning analysis of more than 1800 AHs revealed that ATG3-type AHs contain less hydrophobic and less bulky amino acids compared to unrelated AHs. This biophysical feature of AH_ATG3_ is highly conserved from yeast to mammals, and it is required for ATG3 to achieve efficient LC3 lipidation in cells. Molecular dynamics (MD) simulations of ATG3∼LC3 intermediate showed that the biophysical features of AH_ATG3_ are fine-tuned to regulate its dynamics and substrate accessibility. Moreover, live cell imaging analysis revealed transient membrane association of ATG3 with autophagic membranes, which is governed by the less-hydrophobic feature of AH_ATG3_. Here, ATG3’s AH fine-tuned biophysical features are fundamental to its central role in the ATG3 enzymatic reaction by organizing ATG3∼LC3 conjugate on membranes. We propose that the conceptual and technical framework we present here can serve as a general platform to better understand the role of AHs in regulating key phenomena in the cell.

## Results

### The essential function of AH_ATG3_ for LC3 lipidation is conserved among species

To investigate the sensitivity of AH_ATG3_ to membrane curvature, we first performed circular dichroism (CD) experiments with a peptide corresponding to the N-terminal 22 aa residues of the human ATG3 sequence (ATG3_1-22_ peptide) that contains an amphipathic *α*-helix AH_ATG3_. Consistent with a recent report (*32*), the peptide was unfolded both in aqueous solution and in the presence of extruded DOPC/DOPE liposomes (≧100 nm diameter), while its CD spectrum demonstrated the characteristic shape of an *α*-helix in the presence of sonicated liposomes (∼38 nm diameter) under high protein/lipid ratio (1:80) conditions (**Fig. 1A and B**). We also checked the effect of anionic phospholipids on an *α*-helical conformation of the ATG3_1-22_ peptide by using sonicated liposomes. Compared to DOPC/DOPE liposomes, anionic phospholipids, such as phosphatidylinositol (PI), phosphatidylserine (PS) and phosphatidic acid (PA), slightly promoted the *α*-helical conformation of the ATG3_1-22_ peptide (**Fig. S1B and S1C**). These results indicate that AH_ATG3_ has the ability to interact with highly curved membranes with a preference for charged membranes *in vitro*.

**Figure 1.**
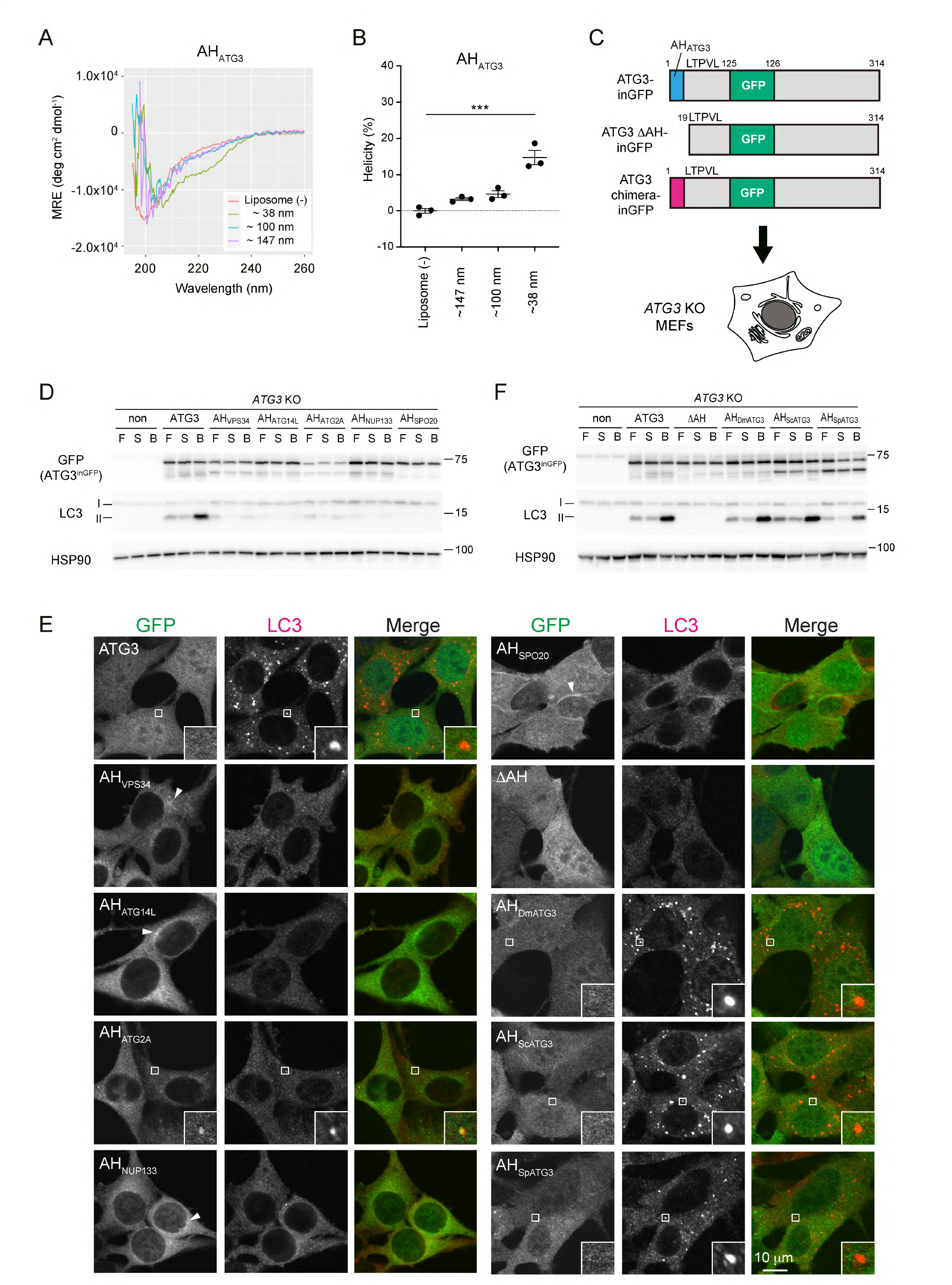
AH_ATG3_ cannot be substituted for by unrelated AHs in human ATG3 protein. (**A**) Far-UV CD spectra of AH_ATG3_ peptide (75 μM) in the absence or presence of DOPC/DOPE (70/30) liposomes (6 mM). A peptide corresponding to the N-terminal ATG3 (1- 22 aa) sequence and containing an additional C-terminal WK residues was used to facilitate titration by UV spectroscopy. The diameter of sonicated liposomes, liposomes extruded through 0.1 μm or 0.2 μm filters were 38 nm, 100 nm, and 147 nm, respectively. MRE, mean residue ellipticity. (**B**) Helicity at 222 nm as determined from the spectra shown in panel **A**. Data represent the mean ± SEM of three independent experiments. (**C**) An experimental scheme of ATG3 rescue assay using ATG3 chimeras. *ATG3 KO* MEFs expressing ATG3 WT, ATG3 ΔAH deletion mutant or ATG3 chimera were used to analyze LC3 lipidation and LC3 puncta formation. The domain organization of ATG3-inGFP is shown: A GFP tag is inserted into the region after E125 residue. P21 and L23 in the conserved region following AH_ATG3_ are coded in all ATG3 constructs. (**D, F**) LC3 flux assay. The indicated cells were starved for 6h with (B) or without 100 nM Bafilomycin A_1_ (S) or cultured in full media (F). Cell lysates were analyzed by immunoblotting using the indicated antibodies. (**E**) LC3 puncta formation. The cells were starved for 1 h, fixed and stained with anti-LC3 antibody. The specimens were analyzed by FV3000 confocal microscope. Arrowheads indicate the membrane localization of ATG3 chimeras. Scale bar, 10 μm. Differences were statistically analyzed by one-way ANOVA and Turkey multiple comparison test. \*\*\**P* < 0.001. See also Figure S1.

To assess whether the membrane binding via AH is sufficient for AH_ATG3_ function *in vivo*, we designed a rescue assay by using ATG3 chimeras in *ATG3* KO MEF cells (**Fig. 1C**). We replaced the AH_ATG3_ region (1-18 aa) with other AHs (*18, 20, 35, 36*) and made various ATG3 chimeras. All constructs kept the key residues P21 and L23 that are critical for LC3 lipidation (*32*) in the conserved region following AH_ATG3_ (**Fig. 1C**). To avoid N- or C-terminal tagging of ATG3 which might interfere with ATG3’s function, we inserted a GFP tag into the middle region of ATG3 (ATG3-inGFP) by following previous work (*37*). To check LC3 flux upon starvation, the cells were cultured under fed or starved conditions with and without lysosomal inhibitor bafilomycin A_1_ (BafA_1_). In this experiment, ATG3-inGFP restored LC3 lipidation under fed conditions and LC3-II accumulation in the presence of BafA_1_, indicating that ATG3-inGFP was fully functional in cells. In contrast, ATG3 chimeras carrying AHs derived from VPS34, ATG14L, ATG2A, NUP133 or Spo20, did not efficiently rescue LC3 lipidation (**Fig. 1D and S1D**). In line with this, LC3 puncta formation upon starvation was restored in the cells expressing ATG3 but not in the cells expressing the ATG3 chimeras (**Fig. 1E and S1E**). Intriguingly, the ATG3 did not show clear punctate structures, while AH_VPS34_, AH_ATG14L_, AH_NUP133_, and AH_Spo20_ chimeras showed a membrane localization and/or puncta formation (**Fig. 1E, arrowheads)**. ATG3 chimera carrying AH_ATG2A_ co-localized with a few small LC3 puncta in some cells, but the puncta size was much smaller than those observed in the cells expressing ATG3 WT. These data indicate that the membrane targeting of ATG3 via any other AH is not enough for ATG3 to efficiently execute LC3 lipidation reaction.

Alternatively, ATG3 chimeras carrying AH derived from other species’ ATG3 clearly restored LC3 lipidation (**Fig. 1F)**. AH_DmATG3_ and AH_ScATG3_, which are AHs of *Drosophila melanogaster* Atg3 and *Saccharomyces cerevisiae* Atg3, respectively, induced LC3 lipidation as much as AH_ATG3_ (**Fig. S1F**). LC3 puncta formation was restored in these cells as well (**Fig. 1E and S1E**). In the cells expressing ATG3 chimera carrying AH_SpATG3_ derived from *Schizosaccharomyces pombe* Atg3, impaired LC3 lipidation and puncta formation were efficiently rescued (**Fig. 1E, 1F, S1E and S1F**). Collectively, these findings suggest that a key function of AH_ATG3_, which cannot be substituted by unrelated AH_non-ATG3_, is conserved in ATG3 proteins.

### Unsupervised machine learning reveals the unique biophysical identity of AH_ATG3_

We searched for common features of AHs derived from ATG3 proteins by comparing sequences of various AHs. ATG3-type AHs have multiple charged and uncharged polar residues and the positively charged residues (K and R) are located at the hydrophilic/hydrophobic interface (**Fig. 2A**). On the other hand, glycine (G) and serine (S) residues are enriched in the hydrophilic faces of AH_ATG14L_ and AH_NUP133_. AH_Spo20_ has a large hydrophilic face composed of many positively charged residues, while AH_ATG2A_ contains many hydrophobic residues. AH_VPS34_ has several charged residues like AH_ATG3_, while it is longer than AH_ATG3_ (**Fig. 2B**). To systematically identify distinct features of AH_ATG3_ conserved among species, we collected homologue sequences of proteins carrying AHs used in the rescue assay (**Fig. 1**), obtained the sequences of their AH region and calculated the amino acid composition and physicochemical properties based on the HeliQuest algorithm (see Materials and Methods) (*38*). Then, we analyzed the dataset of AH parameters using machine learning approaches (**Fig. 2C**). In this analysis, we classified ATG3-type AHs into five groups by phylum: Chordata, Arthropoda, Nematoda, Streptophyta and Ascomycota.

**Figure 2.**
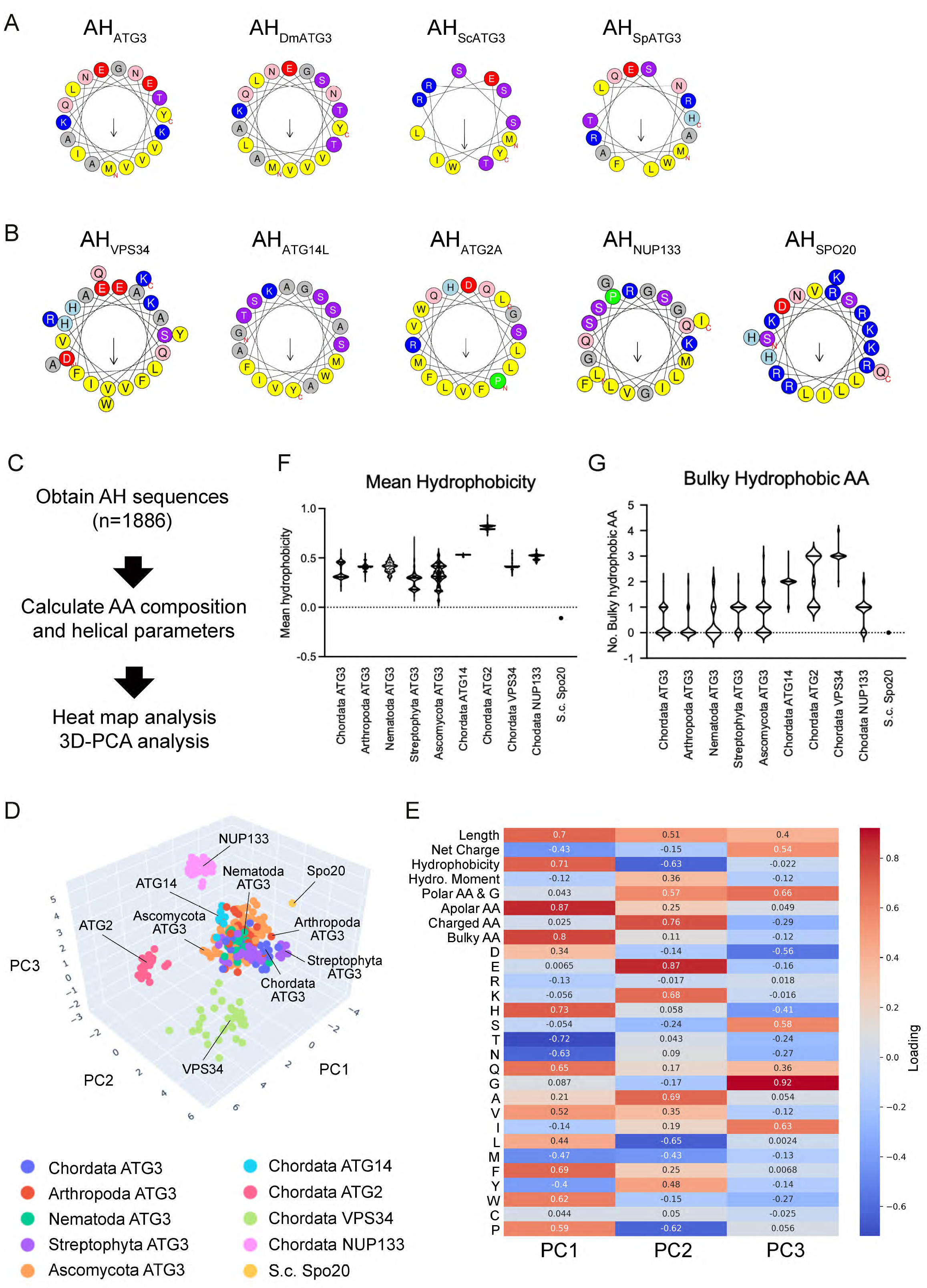
Characterization of conserved AH_ATG3_ features by unsupervised machine learning. (**A, B**) The amphipathic properties of AHs used in the ATG3 rescue assay. Helical wheel representations of AHs derived from ATG3 proteins (**A**) and other unrelated proteins (**B**) were generated in HeliQuest. Hydrophobic residues are shown in yellow, arginine and lysine in dark blue, histidine in light blue, serine and threonine in purple, glutamine and asparagine in pink, proline in green, glutamate and aspartate in red and glycine and alanine in gray. Arrows in helical wheels correspond to the hydrophobic moment. (**C**) Flow chart of AH data analysis. (**D**) Three-dimensional principal component analysis (PCA) of amino acid composition and helical parameter data set. AHs derived from ATG3 proteins were categorized into five groups by phylum: Chordata, Arthropoda, Nematoda, Streptophyta and Ascomycota. Each of the groups is represented by the indicated color. (**E**) PCA Loading matrix. The number of polar residues (Polar AA: S, T, N, H, Q, E, D, K and R), apolar residues (Apolar AA: A, L, V, I, M, Y, W, F, P and C), charged residues (Charged AA: E, D, K and R), and bulky hydrophobic residues (Bulky AA: F and W) were used in the PCA analysis. (**F, G**) Mean hydrophobicity (**F**) and the number of bulky hydrophobic residues (**G**) of AHs. The thick and thin lines in the violin plot represent the medians and quartiles of each group, respectively. See also Figure S2.

We first compared the relative amino acid composition in a heat map (**Fig. S2A**) and noticed that bulky hydrophobic tryptophan (W) residues are rarely found in ATG3-type AHs. Instead, they are rich in small hydrophobic valine (V) residues. Another feature of ATG3-type AHs is a zwitterionic hydrophilic face composed largely of multiple charged and uncharged polar residues. We next investigated ATG3-type AHs by analyzing the dataset of AA composition and AH parameters. To interpret the high-dimensional data, we used 3D- Principal Component Analysis (PCA). PCA efficiently separated ATG3-type AH from AH_ATG2_, AH_VPS34_, AH_NUP133_ and AH_Spo20_ (**Fig. 2D**). The first principal component, PC1, strongly correlates with the number of hydrophobic AA residues, such as the number of apolar residues and bulky hydrophobic residues (**Fig. 2E**). Positive and negative values of PC2 correlate with the number of charged residues and mean hydrophobicity, respectively. PC3 corresponds to the number of glycine (G) residues (**Fig. 2E**). Based on this PCA analysis, we replotted all AHs based on their number of bulky AA and glycine residues, and mean hydrophobicity (**Fig. S2B**). The replotted data showed that ATG3-type AHs are characterized by a lower hydrophobicity, and fewer bulky and glycine residues (**Fig. 2F and S2C**). On the other hand, Chordata AH_ATG14L_ were distributed into the same cluster with ATG3 type AHs in this analysis (**Fig. 2D**). The inability of the ATG3-AH_ATG14L_ chimera to execute LC3 lipidation might be due to the difference in regional effects rather than overall biophysical parameters.

### Low hydrophobicity and a low number of bulky-hydrophobic residues of AH_ATG3_ are crucial for efficient LC3 lipidation

Considering that ATG3 needs to be recruited to membranes to execute LC3 lipidation, the features of AH_ATG3_ —low hydrophobicity and scarce bulky hydrophobic residues— are relatively counterintuitive. Therefore, we investigated whether these AH_ATG3_ features are necessary ATG3’s function in autophagy. We mutated the AH_ATG3_ to introduce hydrophobic residues and performed the LC3 rescue assays. AH_ATG3_ mutants enriched in bulky- hydrophobic residues (3W, 2W and 5W) were prepared by replacing small hydrophobic residues of AH_ATG3_ with bulky tryptophan residues (**Fig. 3A**). The 3W mutant restored LC3 lipidation flux as much as ATG3 WT. In contrast, the 2W and 5W mutants showed less and nearly no LC3 lipidation, respectively (**Fig. 3B and 3C**). LC3 puncta formation was observed in the cells expressing the 3W mutant, whereas smaller punctate structures were formed and the number of LC3 puncta was much less in the cells expressing the 5W mutant (**Fig. 3D and 3E**). These results indicate that increasing both hydrophobicity and the number of bulky residues in AH_ATG3_ is detrimental to ATG3 function *in vivo*. To further validate the significance of the unique biophysical property of AH_ATG3_, we checked the effect of manipulation on the hydrophobic face of AH in the rescue assay. Among AHs used in this study, AH_ATG2A_ shows extremely high hydrophobicity (**Fig. 2G**) and this chimera containing AH_ATG2A_ was not functional in *ATG3* KO cells (**Fig. 1**). The bulky hydrophobic residues (W and F) of AH_ATG2A_ were mutated into valine residues to reduce its hydrophobicity and the number of bulky residues (**Fig. 3A**). In the rescue assay, the AH_ATG3_ chimera carrying less bulky-hydrophobic AH_ATG2A-m1_ significantly restored LC3 lipidation flux and LC3 puncta formation (**Fig. 3D-G**). Thus, the reduction of bulky-hydrophobic residues in the chimeric AH_ATG2_ resulted in a positive impact on ATG3 function.

**Figure 3.**
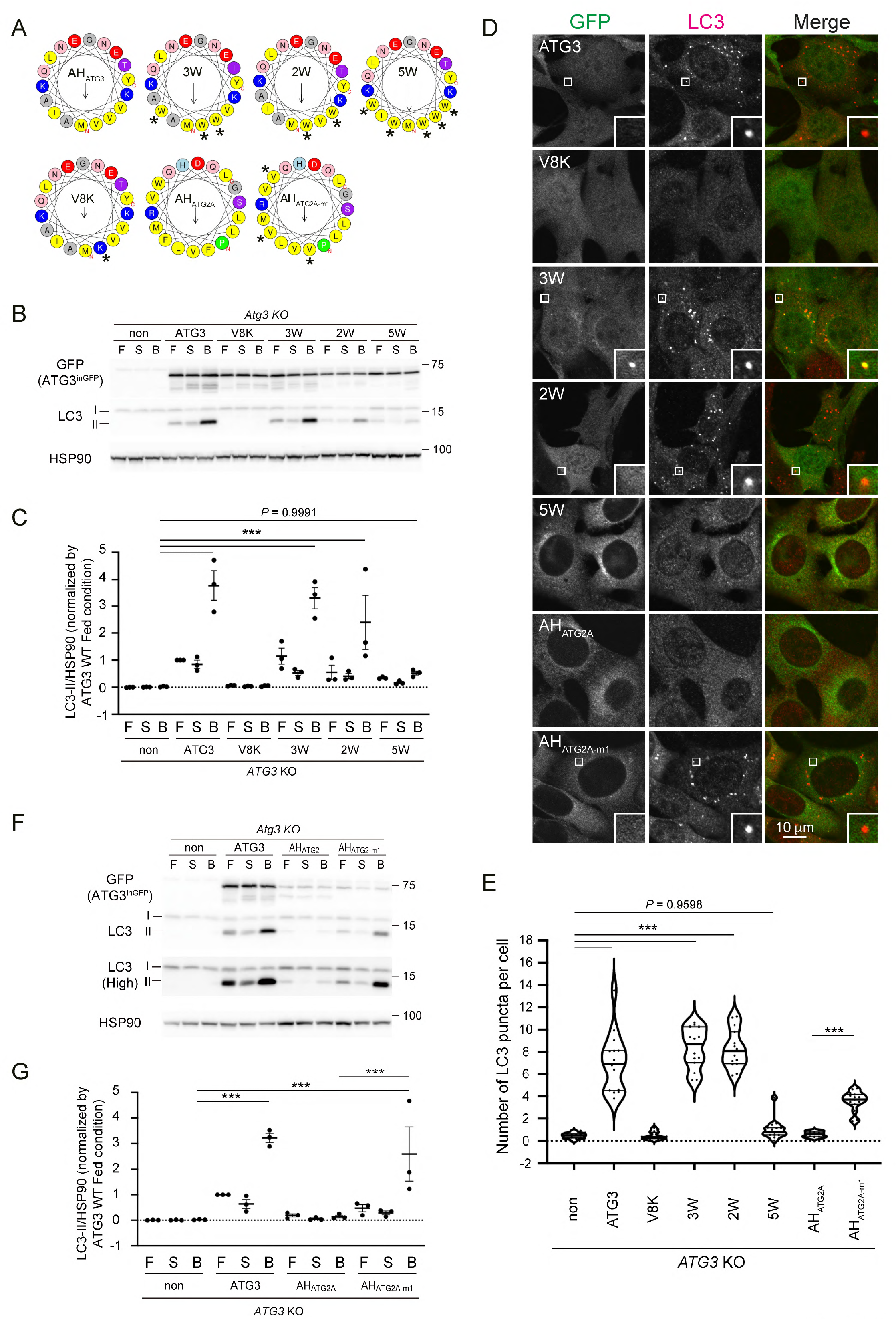
Low hydrophobicity and less bulky features are crucial for AH_ATG3_ to be functional for LC3 lipidation. (**A**) Helical wheel representations of higher bulky-hydrophobic AH_ATG3_ and less bulky- hydrophobic AH_ATG2A_ mutants. V8K mutation destroys the amphipathic character of AH_ATG3_. Asterisks indicate the position of mutations. (**B**) LC3 flux assay of *ATG3* KO cells expressing ATG3 WT or the indicated ATG3 mutants. The cells were starved for 6h with (B) or without 100 nM Bafilomycin A_1_ (S) or cultured in full media (F). Cell lysates were analyzed by immunoblotting using the indicated antibodies. (**C**) Band intensity quantification of LC3-II. All data were normalized with those of HSP90. Data represent the mean ± SEM of three independent experiments. (**D**) LC3 puncta formation. The cells were starved for 1 h, fixed and stained with anti-LC3 antibody. The specimens were analyzed by FV3000 confocal microscope. Scale bar, 10 μm. (**E**) Quantification of the number of LC3 puncta. The thick and thin lines in the violin plot represent the medians and quartiles, respectively. The average number of LC3 puncta per cell was counted from randomly selected areas (n≧14). (**F**) LC3 flux assay of *ATG3* KO cells expressing ATG3 WT or the indicated ATG3 chimeras. Note that ATG3 chimera carrying less bulky-hydrophobic mutant ATG2A-m1 showed the partial restoration of LC3 lipidation. (**G**) Band intensity quantification of LC3-II. All data were normalized with those of HSP90. Data represent the mean ± SEM of three independent experiments. Differences were statistically analyzed by one-way ANOVA and Turkey multiple comparison test. \*\*\**P* < 0.001. See also Figure S3.

Given the relative contribution of the hydrophilic face of AH_ATG3_ on its hydrophobicity, we also examined whether mutations on polar residues affect AH_ATG3_ function (**Fig. S3A**). Replacing either charged (4A mutant) or uncharged polar residues (NTQ-A mutant) did not apparently interfere with LC3 lipidation flux upon starvation (**Fig. S3B and S3C**). Consistently, LC3 puncta formation was clearly observed in the cells expressing the mutants as well as those observed in the expressing ATG3 WT (**Fig. S3D and S3E**). In contrast, the 8A mutant, lacking all polar residues in the hydrophilic face of AH_ATG3_ (**Fig. S3A**), restored neither LC3 lipidation nor LC3 puncta formation (**Fig. S3B-E**), suggesting that overall hydrophilicity (and hydrophobicity as well) of AH_ATG3_ is crucial for ATG3 function rather than some specific polar residues. All together, these data revealed that the defining features of AH_ATG3_ are functionally important in LC3 lipidation reaction.

### MD simulations reveal the dynamics of the ATG3∼LC3 conjugate on the membranes

To study the molecular interaction of ATG3 with membranes during LC3 lipidation reaction, we ran molecular dynamics (MD) simulations and investigated the dynamics of the ATG3∼LC3 conjugate on model membranes. MD simulations produce trajectories that display the dynamics of molecular systems with atomistic details resulting from a physics-based model. In this way, these simulations can provide insight into how dynamics leads to molecular mechanisms of ATG3-dependent LC3–PE conjugation and how these are affected by perturbations of the WT system.

MD simulations require a structure of the complex as initial information. Without an experimentally determined structure, we modeled the complex with AlphaFold (*39*). The five structures predicted by AlphaFold were similar (**Fig. S4A**). ATG3’s N-terminal was still folded as an *α*-helix in these models, connecting via a short linker to the body part of ATG3 containing a large, disordered region. LC3 formed a complex with ATG3 possibly via the LC3 interacting region (LIR) of ATG3 (*40*), and we additionally modeled the thioester bond between CYS264 of ATG3 and GLY120 of LC3. We inserted the Alpha Fold model into a solvated lipid bilayer and ran long equilibrium MD simulations. The simulated complex remained folded and stably inserted into the bilayer.

The ATG3∼LC3 conjugate was highly dynamic and ranged between conformations where LC3 was pointing upwards fully solvated and those bound to the membrane (**Fig. 4A and 4B**). In the initial Alpha Fold model, LC3 pointed away from the membrane, not forming any contact with the membrane (**Fig. 4C**). Its path toward the membrane was sterically hindered by one of ATG3’s disordered regions (125aa-197aa) including the flexible region (89-193aa) (*41–43*) and ATG12-interacting region (153-165aa) (*42, 43*). The disordered region was pointed downwards and charged residues such as GLU170 in the region could form contacts with individual lipids (**Fig. 4C**), tethering the whole region to the bilayer (“LC3- UP/Disordered-DOWN” conformation). ATG3’s N-terminal AH was dynamic throughout all simulations but remained folded and inserted in the membrane (**Fig. S4B**).

**Figure 4.**
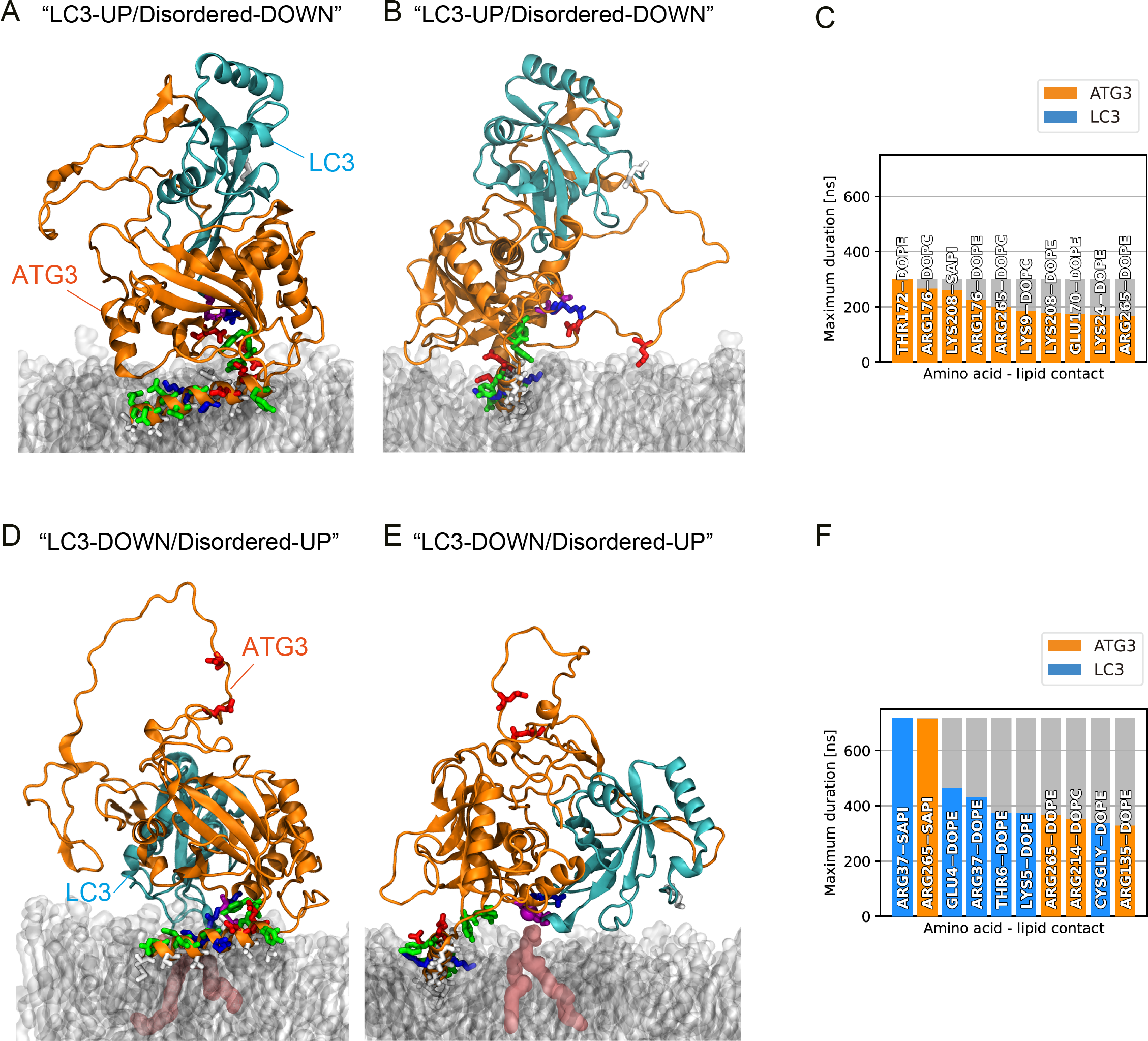
**The ATG3∼LC3 conjugate interacts with the membrane in MD simulations.** (**A**, **B**) Renders of a representative configuration of the ATG3∼LC3 conjugate in the “LC3- UP/Disordered-DOWN” conformation. (**A**) Side view, with the ATG3 N-terminus on the left, and (**B**) axial view, with the terminal part of the AH_ATG3_ closer to the viewer. The renders show the complex in a cartoon representation based on its secondary structure, with ATG3 in orange and LC3 in cyan. The atoms of residues 1-24 (AH and linker), L61, K62, K208, Y209, P263, C264 of ATG3, and M1, K42, F119, G120 of LC3 are also shown explicitly in a licorice representation. The lipid bilayer is shown in transparent gray material. Water and ions are not shown for convenience. (**C**) Histogram of the 10 longest-lasting protein-lipid contacts formed in the “LC3-UP/Disordered-DOWN” conformation. Orange indicates contacts formed between an amino acid of ATG3 and lipid head groups. (**D, E**) Renders of a representative configuration of the ATG3∼LC3 conjugate in the “LC3-DOWN/Disordered-UP” conformation. Rendering and colors as in (**A**, **B**). A PE lipid forming a long-lasting contact with C264 of ATG3 is highlighted in burgundy. (**F**) Histogram of the 10 longest-lasting protein-lipid contacts formed in the “LC3-DOWN/Disordered-UP” conformation. Cyan indicates contacts formed between an amino acid of LC3 and lipid head groups. See also Figure S4.

In our simulations, the complex also spontaneously reorganized to an alternative arrangement, where ATG3’s disordered region pointed upwards and made way for LC3 to approach the bilayer (“LC3-DOWN/Disordered-UP” conformation) (**Fig. 4D and 4E**). In this conformation, LC3 formed long-lasting interactions with lipids in the bilayer. Notably, CYS264 of ATG3 and GLY120 of LC3, connected by the thioester bond, formed a prolonged contact (approx. 400 ns) with a PE headgroup. Additionally, the region around ATG3’s catalytic CYS264 engaged in extended contacts with the lipids (**Fig. 4F**). Most of the lipid interacting residues are relatively conserved (**Fig. S4C, asterisks**), implying that the molecular mechanisms of how ATG3∼LC3 conjugate interacts with membranes might be conserved across species.

### AH_ATG3_ regulates the organization of ATG3∼LC3 conjugate on membranes

We wondered how mutations in the AH would affect its dynamics and the organization of the ATG3∼LC3 conjugate. We created an atomistic model of the first 24 residues of ATG3 as an *α*-helix and inserted it into a lipid bilayer (**Fig. 5A**). Studying the helix in isolation enabled us to focus on the helix-membrane interactions and accumulate longer simulation times. As in the complex, the helix was dynamic but remained inserted and folded during cumulative 30 μs long MD simulations. The helix was localized at the interface between the bilayer’s headgroup region and water. Its hydrophobic residues pointed towards the core of the membrane, whereas the polar and charged residues flanked it or pointed upwards. Flanking residues formed short-lived interactions with lipid head groups. While dynamic, the N-terminus inserted, on average, more deeply into the bilayer than the helix’s C-terminal part. In a control MD simulation, the same *α*-helix model in water rapidly unfolded (**Fig. S5**), confirming that the N-terminal region of ATG3 needs to be inserted into a lipid bilayer to form an amphipathic helix.

**Figure 5.**
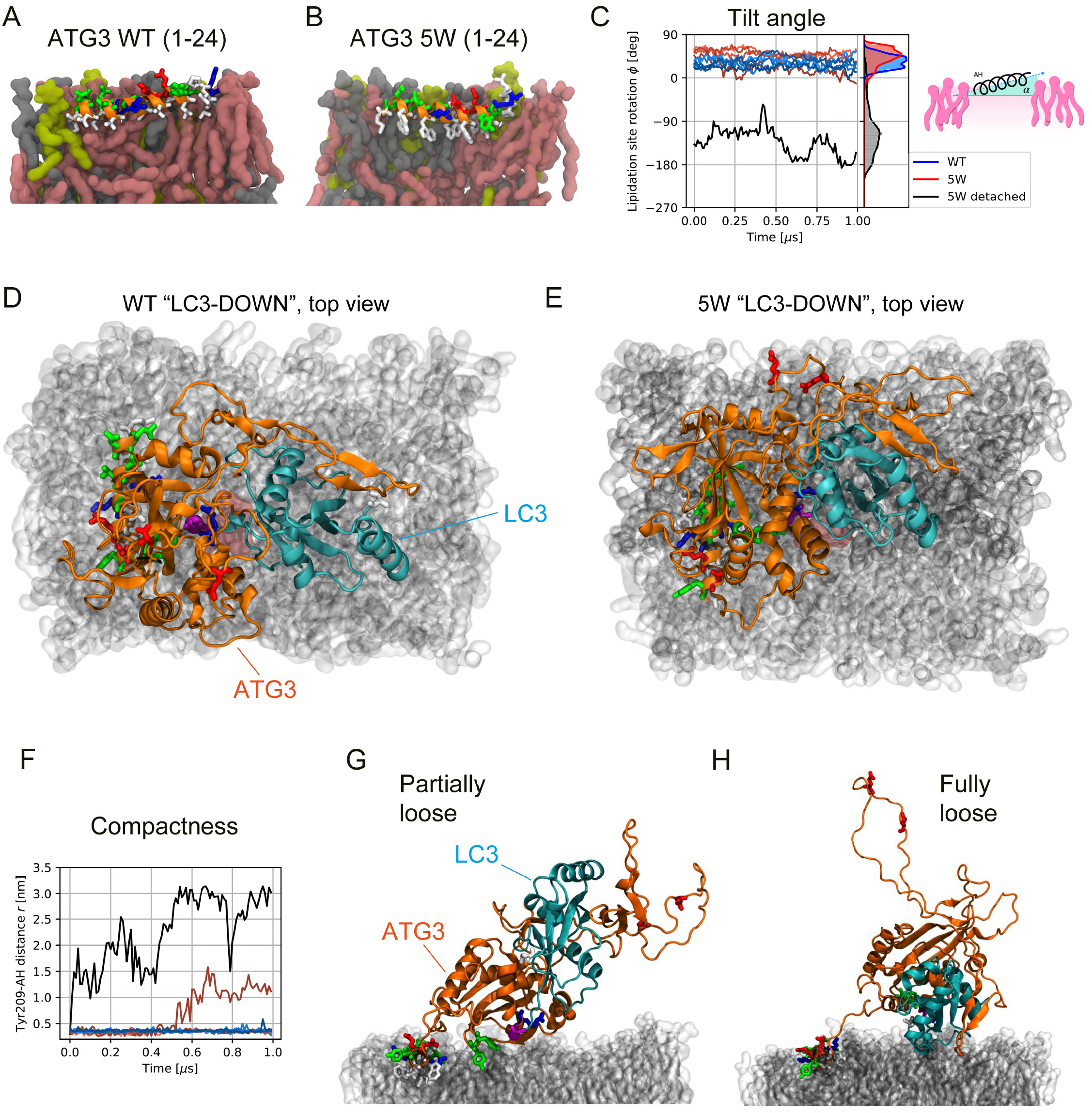
**The 5W AH mutant hinders interactions between LC3 and the membrane.** (**A, B**) Renders of representative molecular configurations of the (**A**) WT AH of ATG3 and (**B**) 5W mutant. The secondary structure and residues of the AH are shown as an orange cartoon and licorice residues, respectively. The renders show a cross-section of the bilayer leaflet where the AHs are inserted. The bilayer contains the same amount of PE, PI, and PC lipids, shown in burgundy, yellow, and gray, respectively. Water and ions are not shown for clarity. (**C**) Time series of the tilt angle alpha formed by the AH in a lipid bilayer. The cartoon inset shows a definition of alpha. Blue and red indicate the values for the WT and 5W mutant, respectively. The right part of the plot reports a histogram of the two title angles. (**D, E**) Top- view renders of two representative configurations of the ATG3∼LC3 conjugate in the “LC3- DOWN” conformation, for (**A**) the WT and (**B**) the 5W mutation. The renders show the complex in a cartoon representation based on its secondary structure, with ATG3 in orange and LC3 in cyan. The atoms of residues 1-24 (AH and linker), L61, K62, K208, Y209, P263, C264 of ATG3, and M1, K42, F119, G120 of LC3 are also shown explicitly in a licorice representation. Both conformations are oriented in the same way, with the AH inserted in the bilayer on the left along the vertical direction from the C-terminus (bottom) to the N-terminus (top). The lipid bilayer is shown in transparent gray material. Water and ions are not shown for convenience. (**F**) Time series of the ATG3∼LC3 conjugate compactness as measured by the distance between TYR209 and the AH. The plot shows 10 time series coming from five simulations of the WT complex and further five simulations of the 5W complex. The black and burgundy curves come from the 5W simulations. Representative conformations adopted by the 5W ATG3∼LC3 conjugate corresponding to the burgundy (**G**) and black (**H**) curves in (**F**), respectively. Rendering and colors as in (**D**, **E**). See also Figure S5.

We considered the atomistic model of the AH inserted in a lipid bilayer and mutated five residues to W (**Fig. 5B**). The 5W mutation, which cannot rescue, remained stably folded and inserted in a different organization in the bilayer during a 3 μs long MD simulation. We quantified the different placements in the bilayer by comparing the distribution of tilt angle alpha of the WT and 5W AHs (**Fig. 5C**). Alpha is the angle formed between the axis of the AH and the XY plane identified by the membrane. For both helices, alpha was highly dynamic over approximately 30 degrees. However, there was a significant difference between WT and 5W: While the former formed, on average, a 5.0 ± .0 deg angle—the helix pointed upwards from the N- to the C-terminus—the latter formed a negative angle, -2 ± 1 deg.

The 5W AH affected the organization of the whole ATG3∼LC3 conjugate and its interactions with the membrane. We simulated the complex containing a 5W mutated AH in the LC3-UP and LC3-DOWN conformations. In the first case, there were no significant differences with the WT-LC3 complex in the analogous conformation. Instead, comparing WT and 5W in the LC3-DOWN conformation showed LC3 and its lipidation site appeared slightly rotated (**Fig. 5D and 5E**). Additionally, the 5W-LC3 complex displayed a more heterogeneous organization. The five simulations of the WT in the LC3-DOWN conformation we ran did not reorganize and remained in contact with the membrane (**Fig. 5F**). On the contrary, out of the five simulations ran with the 5W-LC3-DOWN conformation, two significantly rearranged. The soluble part of the complex lost contacts with the AH decreasing its interactions with the membrane (**Fig. 5G and 5H**).

### Transient membrane association of ATG3 is governed by AH_ATG3_ *in vivo*

We observed that ATG3 WT did not show clear membrane localization in the rescue assay (**Fig. 1E**). Therefore, we hypothesized that the membrane association of ATG3 might be transient. To validate this possibility, we analyzed ATG3-inGFP dynamics in live cells under starved conditions using a Deltavision microscope. As LC3 was unavailable as an autophagosome marker in *ATG3* deficient cells, Halo-tagged ATG5 was used to visualize autophagic membranes instead. Under starved condition with Halo-ligands, Halo-ATG5 puncta gradually appeared and then disappeared within a few minutes in ATG3-inGFP- expressing *ATG3* KO cells (**Movie 1**). On the other hand, ATG3-inGFP was diffusely distributed in the cytoplasm and the fluorescence signal peaks were not clearly observed on Halo-ATG5-positive structures (**Fig. 6A and 6B**). In some cases (∼4.44%), an enrichment of ATG3-inGFP fluorescence signals on Halo-ATG5 puncta was detected (**Fig. 6A, 6B**, **and Movie 2**). We imaged the cells expressing *Δ*AH mutant and analyzed more than 100 Halo- ATG5 puncta in live cells, but there was no correlation of *Δ*AH mutant with Halo-ATG5 (**Fig. 6C, 6F, and Movie 3**), confirming that ATG3-inGFP is targeted to autophagic membranes dependent on its AH_ATG3_. These results suggest that a small amount of ATG3 is sufficient to achieve LC3 lipidation reaction though it cannot be easily visualized by fluorescent microscopy. In contrast, ATG3 chimeras carrying a bulky-hydrophobic rich AH_ATG3_ mutant often formed punctate structures on Halo-ATG5 puncta (**Fig. 6D and 6E**). Both 3W and 5W mutants were recruited to punctate structures synchronously with Halo-ATG5 puncta formation (**Movies 4 and 5**). Live imaging analysis showed that ∼40.9% and ∼80.9% of Halo- ATG5 puncta were co-localized with 3W and 5W mutants, respectively (**Fig. 6F**). In addition, 5W clearly localized in reticular membrane structures and stayed there even after the disappearance of Halo-ATG5 structures (**Movie 5**), indicating that 5W mutant localizes to non-autophagic membranes in addition to autophagic membranes and that the dissociation of ATG3-inGFP 5W from these membranes is significantly delayed. Collectively, these results suggest that ATG3 dynamics is governed by low hydrophobic and less bulky AH, which ensures efficient ATG3 enzymatic reaction and recycling.

**Figure 6.**
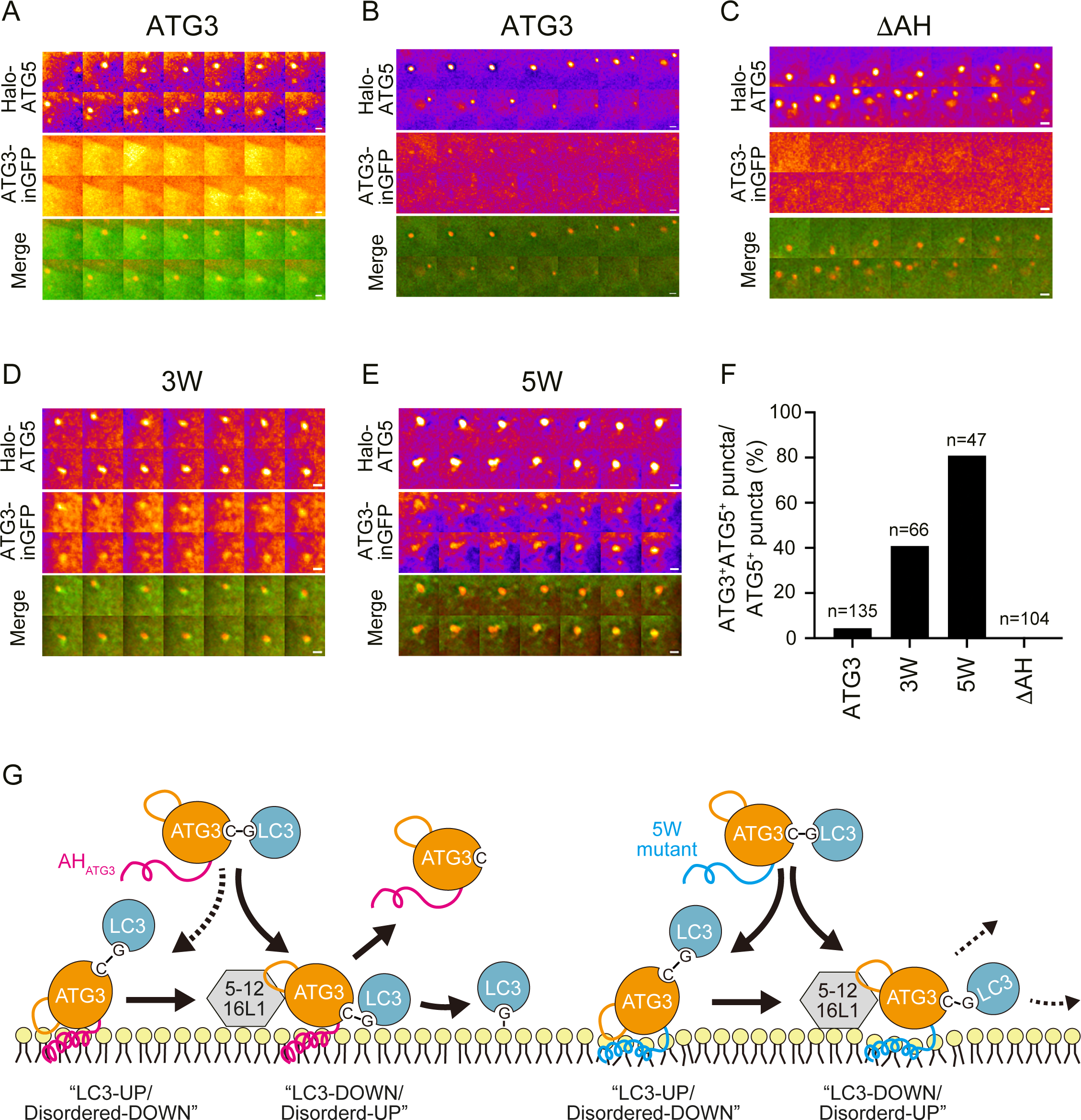
Dynamic behavior of ATG3 on autophagic membranes depends on less bulky-hydrophobic AH. (**A, B**) Live imaging analysis of ATG3-inGFP and Halo-ATG5 under starved condition. The cells were cultured in starvation medium with 200 nM Halo-SaraFluor 650T ligand and observed using a DeltaVision microscopy system. Frames were captured every 30 s. Images are represented in fire lookup table. Two representative images of ATG3 WT are shown. ATG3 was uniformly distributed in the cytoplasm all the time during ATG5 showed a punctate structure (**A**), while an enrichment of ATG3 signal was observed on ATG5 punctum in some cells (**B**). (**C-E**) Representative images of ΔAH deletion mutant (**C**), hydrophobic ATG3 mutants, 3W (**D**) and 5W (**E**). Scale bars, 1 μm. (**F**) Quantification of ATG3- and ATG5-double positive structures in live imaging analysis. (**G**) A speculative model for AH_ATG3_-dependent ATG3 enzymatic reaction. [left] ATG3∼LC3 conjugate is recruited to the membranes via AH_ATG3_. When the disordered region points downwards, LC3 stays away from the membranes (LC3-UP/ Disordered-DOWN). Once the disordered region points upwards and binds to ATG12, the GLY-CYS thioester bond can get access to PE lipids, followed by LC3 lipidation and ATG3 leaving from the membranes (LC3-DOWN/ Disordered-UP). [right] ATG3∼LC3 conjugate carrying 5W mutation in AH_ATG3_ is stably associated with the membranes irrespective of its interaction with ATG12. 5W mutation interferes with the organization and/or stability of ATG3∼LC3 conjugate in the LC3-DOWN conformation, resulting in less LC3 lipidation. See also Movies 1-5.

## Discussion

Our findings converge to support a model of AH_ATG3_-mediated LC3 lipidation reaction on autophagic membranes (**Fig. 6G**). ATG3∼LC3 conjugate can be recruited to autophagic membranes via the membrane association of AH_ATG3_. In this state, LC3 is pointing upwards and does not interact with the membrane due to the steric hindrance by the ATG3 disordered region (**Fig. 4**). Given that ATG3 interacts with ATG12, a component of the E3-like ATG12– 5-16L1 complex, via its ATG12-binding region in the disordered region (**Fig. S4C**) (*42, 43*), a conformational change of ATG3∼LC3 conjugate from “LC3-UP/Disordered-DOWN” to “LC3- DOWN/Disordered-UP” might be facilitated by the interaction of ATG3 with ATG12–5-16L1. While previous data (*41, 42, 44*) has suggested a mechanism for how ATG3 enzymatic activity is promoted by ATG12–5-16L1, our observations support an alternative mechanism. Subsequent to the movement of the disordered region, the CYS-GLY thioester bond of ATG3∼LC3 conjugate gets closer to the lipid head group forming an ATG3∼LC3–PE intermediate. Once LC3 GLY120 forms an amide bond with an amino group of PE, ATG3 promptly dissociates and goes to the next round (**Fig. 6G, left**). This highly dynamic property of ATG3 is determined by the unique biophysical fingerprint of AH_ATG3_ (**Fig. 1-3**). In the case of the high bulky-hydrophobic 5W mutant, ATG3 targets autophagic membranes, is stalled on the membranes and cannot leave efficiently. MD simulation suggests that 5W mutation interferes with the organization and stability of ATG3∼LC3 conjugate in the state of LC3- DOWN conformation (**Fig. 5**), resulting in less lipidation (**Fig. 3B, 3C and 6G, right**). From these findings, we propose that AH_ATG3_ plays a central role in ATG3 enzymatic reaction by organizing the overall structure of ATG3∼LC3 conjugate on membranes and the unique properties of the AH ensure a transient membrane association of ATG3.

Based on the biochemical activity of ATG3 *in vitro* experiments, it has been previously proposed that AH_ATG3_ serves as a curvature sensor in LC3 lipidation reaction (*22*). But, contrary to this assumption, membrane curvature-sensing AHs, such as AH_ATG14L_ and AH_NUP133_, were not functional in the *ATG3* KO rescue assay (**Fig. 1**), indicating that *in vivo* function of AH_ATG3_ is neither just a membrane anchor nor curvature sensing. Then, how does AH_ATG3_ positively contribute to ATG3 enzymatic reaction? One possible scenario is that dynamic interaction of AH_ATG3_ with membranes could allosterically control the structure of ATG3∼LC3 conjugate together with the disordered region (*42*), thus regulating substrate accessibility to the CYS-GLY thioester bond. Another possibility is that AH_ATG3_ might serve as a transient component of the catalytic triad. As we observed that AH_ATG3_ almost reaches both the CYS-GLY thioester bond and PE lipids in the simulation (**Fig. 5F**), AH_ATG3_ might be able to organize the hydrogen bond network with the thioester bond and PE head group during LC3 lipidation reaction, which will promote the deprotonation of PE amino group and the formation of oxyanion hole (**Fig. S1A**). These mechanistic aspects need to be scrutinized in a future study. Recently, it has been reported that ubiquitin can be conjugated to PE (*45*) as well as Atg8/LC3. However, we could not find an AH in the E2 enzyme for PE ubiquitination. As the E3 enzyme involved in the ubiquitin–PE conjugation is a membrane-spanning protein, the ubiquitin transfer cascade for PE lipids might use a distinct mechanism from ATG3- dependent LC3 lipidation reaction.

So far there have been no reported structures for the ATG3∼LC3 conjugate due to the arduous task of the conjugate purification and its intrinsic instability. To overcome this technical issue, we took advantage of AlphaFold model to obtain a predicted structure of ATG3∼LC3 conjugate. To our knowledge, this is the first study to analyze membrane association of the ATG3∼LC3 conjugate *in silico* and show the possibility that its CYS-GLY thioester bond can form a long-lasting contact with lipid head groups. As several residues around the catalytic active site (LYS208, TYR210, ARG214, and ARG265) show the interaction with lipids within a narrow region together with the CYS-GLY thioester bond, these residues could be at play in the enzymatic reaction. In addition, the disordered region (ARG135, GLU170) of ATG3 and the N-terminal region (LYS5) of LC3 are associated with the membranes. Consistently, the lipidated LC3 N-terminus has been shown to interact with liposomes *in vitro* and *in silico* (*14*). These weak membrane associations might work in a concerted manner, in tandem with the ATG16L1 and WIPI2 proteins (*33*), to facilitate LC3 lipidation and/or autophagosome biogenesis.

Accumulating evidence shows that some other ATG proteins have functional AHs. The requirement for an amphipathic region and the dependency on membrane curvature *in vitro* have been investigated in previous studies. However, it is still enigmatic how AH_ATG_ contribute to autophagy *in vivo*, partly because our knowledge on the physicochemical properties of AH_ATG_ essential for autophagy is very limited. Indeed, the gap between *in vitro* and *in vivo* experiments makes it difficult to assess AH_ATG_ biological functions. In this study, we revealed that AH_ATG3_ are characterized as being less bulky-hydrophobic compared to other AHs, and this property is essential for its *in vivo* function. Given that bulky hydrophobic residues increase the membrane penetration activity of AHs, membrane association mediated by AH_ATG3_ might be energetically advantageous for the repetitive lipidation reactions required to decorate the membrane with lipidated ATG8 proteins. It would be important, and indeed as our work shows, essential to investigate the dedicated function of each AH_ATG_ based on their unique properties of each AH_ATG_ to deepen the understanding of the relationship between ATG proteins and membrane lipids.

Fine-tuning the relationship between structure at the atomistic resolution, membrane lipid composition, and dynamics could be a general feature for AHs involved in cellular processes (*46, 47*). We propose to move beyond simplified models, which are based exclusively on one biophysical property, to a more nuanced understanding of the role played by AHs. From this point of view, comparative analysis of AHs with unsupervised machine learning will be a powerful approach for the identification of multiple key properties conserved among species. Moreover, we showed that the dynamics of the AH_ATG3_ in the membrane controls the structural dynamics of the whole complex. The perturbation induced by mutating amino acids in AH_ATG3_ could also be mimicked by changing the composition of the lipids that surround the helix. AHs and their host membranes co-evolved to enable fundamental cellular functions and should be studied with a holistic approach (*48*).

## Materials and Methods

### Plasmids

Descriptions of plasmids used in this study are provided in Table S1. cDNAs encoding the full length of human ATG3 (NP_071933) and mouse ATG5 (NP_444299) were amplified by PCR and subcloned into pMRX-IP and pMRX-IPU-HaloTag7 backbone vectors, respectively. pMRX-IP-hATG3_1-125_-EGFP-hATG3_126-314_ was generated by inserting the sequence coding EGFP into between E125 and I126 of ATG3. For generation of ATG3 chimeras carrying AH derived from other proteins, two overlapping oligonucleotide primers coding AH of interest were annealed and elongated by a single PCR reaction. Subsequently, the purified PCR products were inserted by seamless DNA cloning method (*49*). ySpo2057-77aa (NP_013730), hNUP133_245-265aa_ (NP_060700), hVPS34_864-867aa_ (NP_002638), hATG14L_471-488aa_ (NP_055739), hATG2A_1750-1767aa_ (NP_055919), *Drosophila melanogaster* Atg3_1-18aa_ (NP_649059), *Saccharomyces cerevisiae* Atg3_1-12aa_ (NP_014404), and *Schizosaccharomyces pombe* Atg3_1-15aa_ (NP_596664) were described as AH_Spo20_, AH_NUP133_, AH_VPS34_, AH_ATG14L_, AH_ATG2A_, AH_DmATG3_, AH_ScATG3_, and AH_SpATG3_, respectively. All deletion and point mutation mutants of ATG3 were generated by a PCR-based method.

### Antibodies and reagents

Antibodies used for immunoblotting are listed: rabbit polyclonal anti-LC3 (#1) (*50*), mouse monoclonal anti-HSP90 (BD Transduction Laboratories, 610419), rabbit polyclonal anti-GFP (Invitrogen, A6455), Secondary antibodies are HRP-conjugated anti-rabbit IgG (Jackson ImmunoResearch Laboratories, 111-035-144), HRP-conjugated anti-mouse IgG (Jackson ImmunoResearch Laboratories, 315-035-003). Antibodies used for immunofluorescence are as listed: rabbit polyclonal anti-LC3 (MBL, PM036, lot 025) and AlexaFluor 568-conjugated anti-rabbit IgG (Invitrogen, A-11036). All lipids were purchased from Avanti: DOPC (850375C), DOPE (850725C), Liver PI (840042C), DOPS (840035C) and DOPA (840875C). Bafilomycin A_1_ (#11038) was purchased from Cayman.

### Circular dichroism (CD) spectroscopy

The peptide containing ATG3 N-terminal region (MQNVINTVKGKALEVAEYLTPVWK) was synthesized by the Francis Crick Peptide Chemistry Technology Platform. Lipids were mixed at the desired ratio, dried under argon and placed in a rotary evaporator to completely remove the organic solvent. The lipid films were rehydrated and resuspended in buffer A (10mM Tris-HCl pH 7.5, 150mM KCl). The suspension was frozen and thawed 5 times using liquid nitrogen and a water bath, and then extruded through 0.2 μm or 0.1 μm membrane

(Whatman) using a Mini-Extruder (Avanti Polar Lipid). Very small liposomes were prepared by sonication on ice with a tip sonicator (Misonix). Lipid debris were removed by 25,000 x*g* for 20 min. The size of liposomes was checked by Zetasizer Nano ZS (Malvern Instruments). For CD spectra measurement, the ATG3 peptide (final conc. 75 μM) was mixed with 6 mM liposome solution and analyzed using a Jasco J-815 spectrometer with a quartz cell of 0.02 cm path length. Each spectrum is the average of 50 scans recorded from 190 to 260 nm, with a bandwidth of 2 nm, a step size of 0.2 nm and a scan speed of 200 nm min^-1^. The buffer contribution was subtracted. The *α*-helix content was calculated from the molar ellipticity at 222 nm ([∂]_222 nm_) according to: % *α*-helix ([∂]_222 nm_+2340)/303 (*36*).

### Cell lines and culture conditions

Authenticated HEK293T cells were used in this study. *Atg3* KO MEF was generated previously (*51*). Cells were maintained in Dulbecco’s Modified Eagle Medium (Wako, 043- 30085) supplemented with 10% fetal bovine serum (Sigma-Aldrich, 173012) in a 5% CO_2_ incubator at 37 °C. *Atg3* KO MEF stably expressing ATG3 WT or ATG3 chimera were generated as follows: HEK293T cells were transfected using Lipofectamine 2000 reagent (Thermo Fisher Scientific, 11668019) with retroviral plasmid, pCG-VSV-G and pCG-gag-pol, following which the medium was replaced with fresh medium. After 3 days, the culture medium was collected and filtered with a 0.45 μm filter unit (Millipore, SLHVR33RB). *Atg3* KO MEFs were treated with the retrovirus containing medium and 8 μg/mL polybrene (Sigma-Aldrich, H9268). After 2 days, drug selection was performed with 3 μg/mL puromycin (Sigma-Aldrich, P8833). Subsequently, GFP-low expressing cells were sorted by flow cytometry (Sony, SH800).

### Immunoblotting

Cells were cultured under DMEM supplemented with FBS or DMEM without amino acids (Wako, 048-33575) in the absence or presence of 100 nM bafilomycin A_1_ for 6 h, and then collected in ice-cold PBS by scraping on ice. The precipitated cells were suspended in 100 μl lysis buffer (25 mM Hepes-NaOH, pH7.5, 150 mM NaCl, 2 mM MgCl_2_, 0.2% n-dodecyl-b-D-maltoside [nacalai, 14239-54] and protease inhibitor cocktail [nacalai, 03969-34]) and incubated on ice for 20 min. Ninety μl of cell lysates were mixed with 10 μl of lysis buffer containing 0.1 μl benzonase (Merck Millipore, 70664) and further incubated on ice for 15 min. The remaining cell lysates were centrifuged at 17,700 x *g* for 15 min and the supernatant was used to measure protein concentration by NanoDrop One spectrophotometer (Thermo Fisher Scientific). The cell lysates were mixed with SDS-PAGE sample buffer and heated at 95°C for 5 min. Samples were subsequently separated by SDS-PAGE and transferred to Immobilon-P PVDF membranes (Merck Millipore, IPVH00010) with Trans-Blot Turbo Transfer System (Bio-Rad). After incubation with the indicated antibodies, the signals from incubation with SuperSignal West Pico PLUS Chemiluminescent Substrate (Thermo Fisher Scientific, 34580) was detected with Fusion Solo S (VILBER). Band intensities were quantified with Fiji.

### Fluorescence microscopy

Cells grown on coverslips were fixed with 4% paraformaldehyde in PBS for 20 min, permeabilized with 50 μg/ml digitonin (Sigma-Aldrich, D141) in PBS for 5 min, blocked with 3% BSA in PBS for 30 min. After incubation with the indicated antibodies, the specimens were mounted in SlowFade with DAPI (Invitrogen, S36939) and observed using a confocal FV3000 confocal laser microscope system (Olympus). For the final output, images were processed using Adobe Photoshop 2021 v22.5.9 software (Adobe). The number of LC3 puncta was counted using Fiji. Briefly, the images were applied with a Gaussian filter for noise suppression and processed with Top-Hat filter. After setting threshold and watershed segmentation, the number of LC3 puncta were counted by Analyze Particles command. Cells were counted by using Cell Counter plug-in. Microcopy and image analysis were done blind.

### Live imaging analysis

*ATG3* KO MEFs stably expressing ATG3-inGFP, mRuby3-LC3 and Halo-ATG5 were grown on a glass bottom dish (IWAKI, 3910-035). The cells were washed twice with PBS and incubated in starvation medium with 200 nM SaraFluor 650T ligand. Images were acquired with a 60× PlanAPO oil-immersion objective lens (1.42 NA; Olympus), using a Deltavision microscope system (GE Healthcare) coupled with an Olympus IX81-ZDC and a cooled CCD camera CoolSNAP HQ2 (Photometrics, Tucson). During live-cell imaging, the culture dish was mounted in a chamber INUB-ONI-F2 (TOKAI HIT) to maintain the culture conditions (37°C and 5% CO_2_). ATG3-inGFP, mRuby3-LC3B and Halo-ATG5 were illuminated with a mercury arc lamp attenuated to 10, 32 and 32% by neutral density filters, respectively. Exposure times were 0.6 s for ATG3-inGFP, 0.6 s for mRuby3-LC3B and 0.4 s for Halo-ATG5. Time-lapse images were acquired at 30-s intervals. Deconvoluted images were obtained using softWoRx software. To analyze ATG3 puncta formation on autophagic membranes, the area containing ATG5-positive puncta in time-series image stacks were selected, converted to an intensity map using the Fire lookup table and then aligned into a single montage image using Fiji. For the final output, images were processed using ImageJ and Adobe Photoshop 7.0.1 software.

### AH data analysis

We collected 1886 protein sequences containing an AH across different species in HMMER web server. Homologue sequences were obtained by using hATG3 (NP_071933), hATG14L (NP_055739), hATG2A (NP_055919), hVPS34 (NP_002638) and hNUP133 (NP_060700) used as a reference sequence. We grouped the sequences by the family and phylum. The data set included 195 Chordata ATG3, 129 Arthropoda ATG3, 44 Nematoda ATG3, 177 Streptophyta ATG3, 419 Ascomycota ATG3, 157 Chordata ATG14, 366 Chordata ATG2, 219 Chordata VPS34, and 179 Chordata NUP133 and one yeast Spo20 protein sequences. The sequences were aligned on MEGA X software and the AH region was extracted. We calculated the amino acid composition and the physicochemical properties of the AHs based on HeliQuest algorithm (*38*). To analyze a large data set, we used a Python code “Hydrophobic Moment Calculator” (*52*) on GitHub with slight modifications. We computed the length, net charge, hydrophobicity, hydrophobic moment, the number of polar (S, T, N, H, Q, E, D, K and R), apolar (A, L, V, I, M, Y, W, F, P and C), charged (E, D, K and R), and bulky hydrophobic (F and W) residues, and the count of amino acids by residue type, for each AH in the data set. We normalized each feature by subtracting the mean and dividing by the standard deviation and analyzed the data with a custom-written Python code based on the Scikit-learn library (*53*). Subsequently, we conducted the principal component analysis (PCA) and restricted to the first 3 components, as they explain more than 50% of the data’s variance. The importance of each feature within a principal component was evaluated according to the loading matrix.

### Molecular modelling and system preparation

We obtained the human ATG3 and LC3B amino acid sequences from UniProt (entries Q9NT62 and Q9GZQ8, respectively (*54*)); of the latter, we selected residues 1-120. We ran two sets of atomistic molecular dynamics (MD) simulations. First, we simulated residues 1- 24 of ATG3 (sequence MQNVINTVKGKALEVAEYLTPVLK) modeled as an alpha helix with UCSF Chimera (*55*). Second, we simulated the entire ATG3∼LC3 conjugate. In the absence of an experimental structure, we modeled it with AlphaFold (*39, 56*). We entered the sequences and used the MMseqs2 homology search and pLDDT confidence measure to rank the models (*57*). We resolved 5 heterodimeric structures and selected the one with LC3 closest to the lipid bilayer.

We used CHARMM-GUI (*58, 59*) to model the lipid membrane and horizontally insert the N-terminus of ATG3 in the upper leaflet (Membrane Builder tool (*60*)) and to add the solvent and ions. In half of the systems, we point-mutated the ATG3 sequence. Finally, we created the topology and structure files for the MD simulations (*61*). The two versions of ATG3 are the wild-type (WT) and the 5W-mutated. In the latter, we turn Val4, Val8, Ala12, Val15, and Ala16 into Tryptophan to increase the N-terminal helix’s hydrophobicity.

We modeled a lipid bilayer with 55% 18:1/18:1-dioleoylphosphatidylethanolamine (DOPE), 30% 18:1/18:1-dioleoylphosphatidylcholine (DOPC), and 15% 18:0/20:4-1-stearoyl-2- arachidonoyl phosphatidylinositol (PI) with the CHARMMGUI membrane builder (*59, 60*). We placed the membrane leaflets horizontally in the simulation box. We inserted the ATG3’s N-terminus parallel to the upper leaflet, with the helix’s hydrophobic face pointing down. We capped the N and C-termini in all peptides to make them neutral. We used TIP3P water solvent and neutralized the system’s net charge by adding 0.18 M of NaCl ions. We imposed periodic boundary conditions on the simulation boxes.

The systems in the first set have approx. 60k atoms and 8 × 8 × 9.8 nm^3^ boxes. Those in the second have 160k atoms and 10 × 10 × 15.8 nm^3^ boxes.

### MD Simulations

We used the CHARMM36m force field (*61, 62*), and ran the simulations with GROMACS 2021.4 (*63–69*). We followed the energy minimization and 1 ns equilibration protocol provided by CHARMM-GUI. We launched NPT production runs with temperature *T* = 310 K (Nose-Hoover thermostat (*70*)), pressure *P* = 1 atm (semi-isotropic Parrinello-Rahman barostat (*71*)), and a 1.2 nm cutoff for both the Van der Walls and the Coulomb forces (particle mesh Ewald method (*72*)). We used a *dt* = 2 fs integration step and saved the configurations to GROMACS xtc trajectory files every 1 ns.

We modeled the thioester bond between Cys264 of Atg3 and Gly120 of LC3 as an “intramolecular interaction” in the GROMACS topology file. We applied a harmonic restraint between the Cysteine’s sulfur and one of the Glycine’s terminal oxygens after the original equilibration, with target distance *d* = 0.12 nm and force constant *k* = 6000 kJ · mol^−1^nm^−2^. Lower *d* and higher *k* resulted in simulation instabilities. We ran further 500 ps before starting the production run; the linked atoms ended up about 0.21 nm distant.

We describe the simulations in the following table:

**Table 1.** Simulated systems Replicas wt thioester run2, run7, run8, run9, and run10, 5w thioester run2, run7, run8, run9,and run10, and wt run4 were initiated from the 5w, run1 configuration at t = 4.7*μ*s. After careful inspection, we took such configuration as representative of the “down” conformation. In the WT simulations, we mutated it back to the original amino acid sequence with CHARMM-GUI.

| Name | Description | Replica | Duration | Notes |
| --- | --- | --- | --- | --- |
| wt ah | Wild-type residues 1-24 | run1 | 6.131 $\mu\text{s}$ | |
| | of Atg3 on the lipid | run2 | 5.036 $\mu\text{s}$ | |
| | membrane | run3 | 5.016 $\mu\text{s}$ | |
| 5w ah | 5W-mutated residue 1-24 | run1 | 3.113 $\mu\text{s}$ | |
|  | of Atg3 on the lipid |  |  |  |
|  | membrane |  |  |  |
| wt thioester | Wild-type Atg3-LC3 with the lipid membrane and the Cys-Gly thioester bond | run1 | 6.228 μs | Initiated from the Alphafold “up” configuration. |
| | | run3 | 0.146 μs | |
| | | run4 | 0.146 μs | |
| | | run5 | 0.146 μs | |
| | | run6 | 0.147 μs | |
| | | run2 | 6.060 μs | Initiated from the “down” configuration of 5w, run1 at $t = 4.7 \mu$s, mutated back to WT. |
| | | run7 | 0.437 μs | |
| | | run8 | 0.435 μs | |
| | | run9 | 0.297 μs | |
| | | run10 | 0.299 μs | |
| 5w thioester | 5W-mutated Atg3-LC3 with the lipid membrane and the Cys-Gly thioester bond | run1 | 6.231 μs | Initiated from the 5W-mutated Alphafold “up” configuration. |
| | | run3 | 0.146 μs | |
| | | run4 | 0.146 μs | |
| | | run5 | 0.146 μs | |
| | | run6 | 0.148 μs | |
| | | run2 | 4.843 μs | Initiated from the “down” configuration of 5w, run1 at $t = 4.7 \mu$s. |
| | | run7 | 0.450 μs | |
| | | run8 | 0.450 μs | |
| | | run9 | 0.316 μs | |
| | | run10 | 0.316 μs | |
| wt | Wild-type Atg3-LC3 with the lipid membrane without the Cys-Gly thioester bond | run1 | 12.282 μs | Initiated from the Alphafold “up” configuration. |
| | | run2 | 8.108 μs | |
| | | run3 | 7.963 μs | |
| | | run4 | 7.462 μs | Initiated from the “down” configuration of 5w, run1 at $t = 4.7 \mu$s, mutated back to WT. |
| 5w | 5W-mutated Atg3-LC3 with the lipid membrane without the Cys-Gly thioester bond | run1 | 12.225 μs | Initiated from the 5W-mutated Alphafold “up” configuration. |
| | | run2 | 9.129 μs | |
| | | run3 | 9.042 μs | |

### MD Trajectory analysis

We used the GROMACS trjconv tool to process the trajectories, and VMD 1.9 (*73*) for visual inspection and rendering. We kept the membrane leaflets horizontal and aligned the central axis of the AH’s C*α* to the **x**^ direction once projected to the *XY* plane (N to C terminus). We performed numerical analysis in custom-written Python code based on the NumPy, SciPy, and MDTraj libraries (*74*).

We described the systems’ dynamics with the following features: *α*: the tilt angle of the central axis of the AH with respect to the horizontal plane; *r*: the minimum distance between Atg3’s TYR209 and the AH heavy atoms —it is a measure of the “compactness” of the complex; We quantified the amino acid-lipid contacts by looking at the minimum distances *d* between the residue’s heavy atoms and lipid phosphate heads. We considered the contact formed for *d <* 0.45. We computed the maximum persistence time for each contact as its most extended continuous series in a simulation. We enforced comparability by considering subtrajectories of equal length either in the “up” or “down” configurations (wt thioesterrun1 and run2), first 6 *µ*s).

### Statistics and reproducibility

Differences were statistically analyzed by one-way ANOVA and Turkey multiple comparison test. Statistical analysis was carried out using GraphPad Prism 9 (GraphPad Software). All data are presented as the mean±SEM. Reproducibility of all results reported here was confirmed.

## Acknowledgements

We thank Keiko Karasawa for the technical assistance with immunoblotting experiments, Simone Kunzelmann for the technical assistance with CD spectra analysis, Shoji Yamaoka for pMRXIP, Teruhito Yasui for pCG-VSV-G and pCG-gag-pol and João Rodrigues for the script of Hydrophobic Moment Calculator. We thank Masaaki Komatsu for providing *Atg3* KO MEFs. This study was supported by PRESTO (JPMJPR20EC to T.N.) and ERATO (JPMJER1702 to N.M.) from Japan Science and Technology (JST), a Grant-in-Aid for Transformative Research Areas (B) (grant 21H05146 to T.N.) from the Japan Society for the Promotion of Science (JSPS), and a grant from the Japan Foundation for Applied Enzymology (to T.N.). S.A.T, was supported by The Francis Crick Institute which receives its core funding from Cancer Research UK (CC2134, CC2064), the UK Medical Research Council (CC2134, CC2064). This research was funded in whole, or in part, by the Wellcome Trust (CC2134, CC2064). G.L. and R.C. acknowledge the support of the Frankfurt Institute of Advanced Studies, the LOEWE Center for Multiscale Modelling in Life Sciences of the state of Hesse, the CRC 1507: Membrane-associated Protein Assemblies, Machineries, and Supercomplexes, and computational resources and support by the Center for Scientific Computing of the Goethe University and the Jülich Supercomputing Centre. R.C. acknowledges the support of the International Max Planck Research School on Cellular Biophysics. For the purpose of Open Access, the author has applied a CC-BY license to any Author Accepted Manuscript version arising from this submission. All data needed to evaluate the conclusions in the paper are present in the paper and/or the Supplementary Materials. The molecular dynamics simulations are publicly available at the repository DOI: 10.5281/zenodo.7614455.

## Author Contributions

T.N. and S.A.T. conceived the project. T.N., R.C., and S.A.T. designed the research. T.N., G.L., and R.C. performed most experimental work and analyzed the data. T.N. and R.C. performed the AH data analysis. G.L. and R.C. designed, performed, and analyzed the MD simulation. T.N. performed the live imaging analysis. T.N., G.L., N.M., R.C., and S.A.T. wrote the manuscript with input from all authors.

---

**Figure S1.**
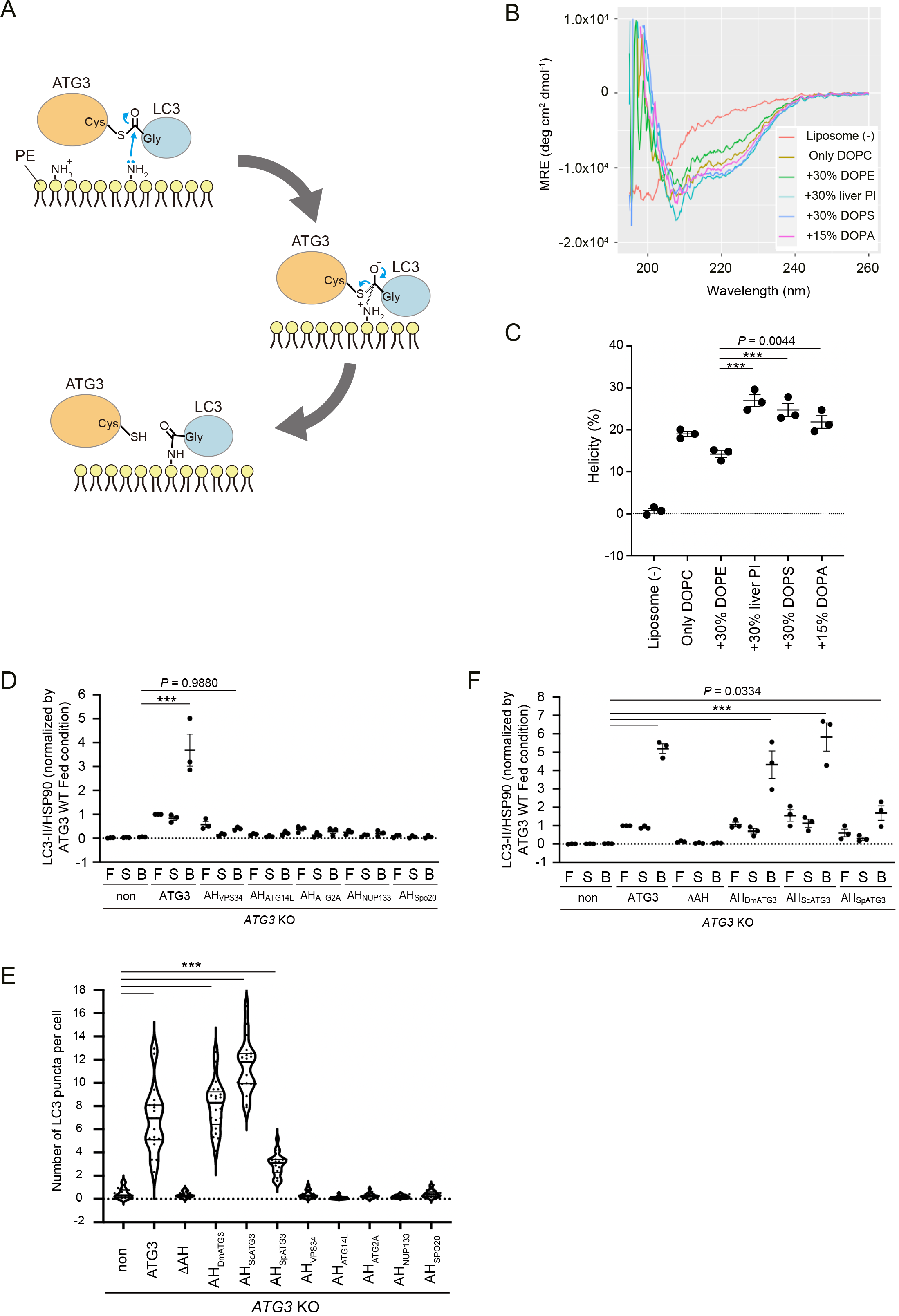
related to Figure 1. **ATG3-dependent LC3 conjugation reaction and quantification results of ATG3 rescue assay.** (**A**) The chemical mechanism of LC3–PE conjugation catalyzed by ATG3. (top) An ATG3∼LC3 conjugate interacts with membranes. The primary amine of PE head group, which is mostly in the protonated state at physiological pH, becomes deprotonated for nucleophilic attack on the thioester bond linking ATG3 and LC3. (middle) After nucleophilic attack, the negatively charged thioester oxygen atom is formed, and it is expected to be stabilized by surrounding residues. (bottom) Simultaneously, ATG3 is dissociated from the intermediate structure and leaves the membranes, which results in the formation of an amide bond between LC3 and PE. (**B**) Far-UV CD spectra of AH_ATG3_ peptide (75 μM) in the absence or presence of sonicated liposomes (6 mM) containing increasing mol% of either DOPE, liver PI, DOPS or DOPA. The remaining lipids in the liposome were DOPC (the concentration of which varied from 70 mol% to 100 mol% depending on the concentration of another lipid). MRE, mean residue ellipticity. (**C**) Helicity at 222 nm as determined from the spectra shown in Fig. S1B. (**D**) Band intensity quantification of LC3-II shown in Fig. 1D. All data were normalized with those of HSP90. (**E**) Quantification of the number of LC3 puncta shown in Fig. 1E. The thick and thin lines in the violin plot represent the medians and quartiles, respectively. The average number of LC3 puncta per cell was counted from randomly selected areas (n≧19). (**F**) Band intensity quantification of LC3-II shown in Fig. 1F. All data were normalized with those of HSP90. Data represent the mean ± SEM of three independent experiments (**C, D, F**). Differences were statistically analyzed by one-way ANOVA and Turkey multiple comparison test. \*\*\**P* < 0.001.

**Figure S2.**
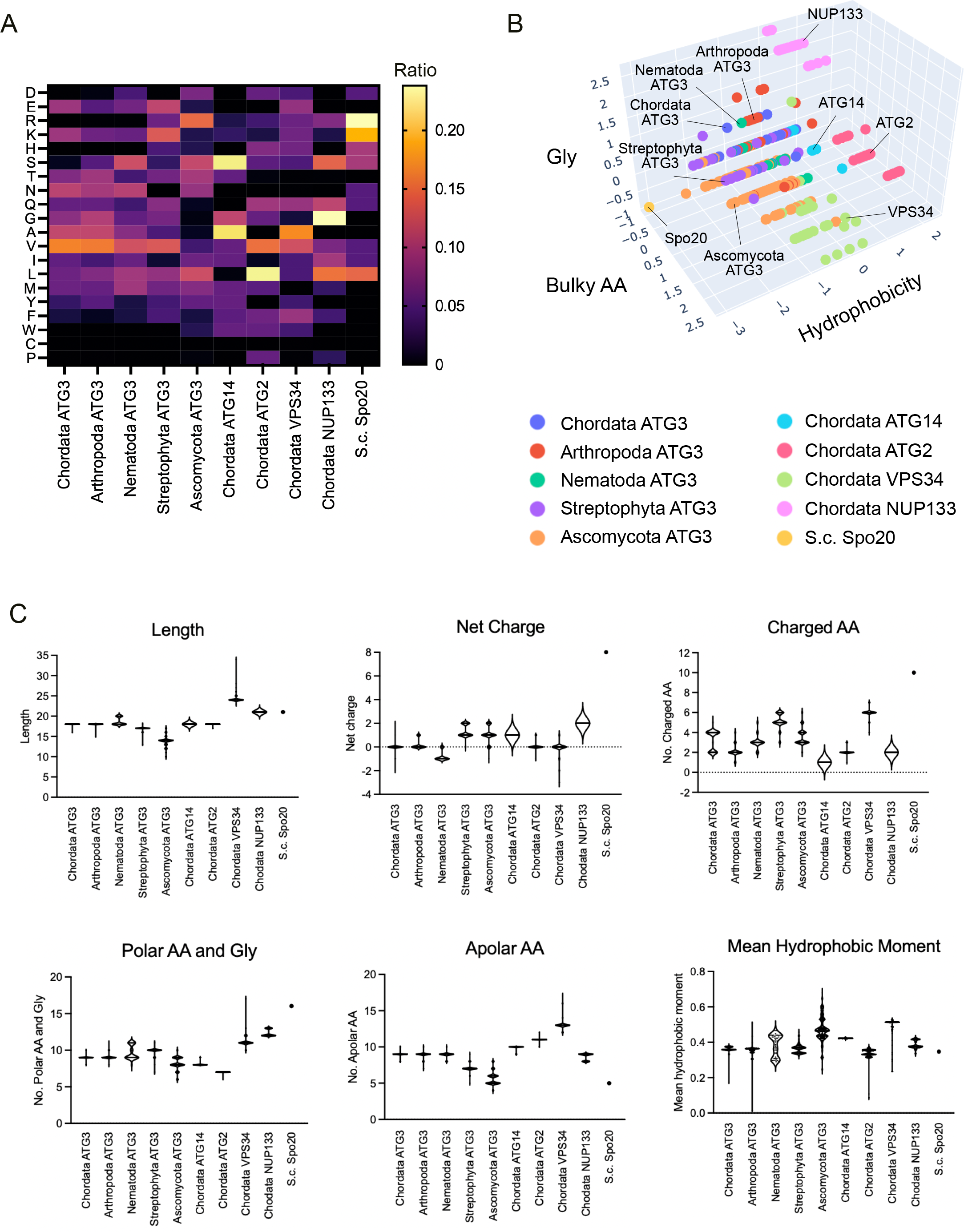
related to Figure 2. Helical parameters of AHs analyzed in this study. (**A**) Heatmap of amino acid composition of AHs. AHs derived from ATG3 proteins were categorized into five groups by phylum: Chordata, Arthropoda, Nematoda, Streptophyta and Ascomycota. (**B**) A 3D-plot of the number of bulky residues, glycine residues and mean hydrophobicity of AHs analyzed in this study. Each of the groups is represented by the indicated colors. (**C**) Length, net charge, the number of charged residue (Charged AA), the number of polar residue and glycine residue (Polar AA and Gly), the number of apolar residue (Apolar AA) and mean hydrophobic moment are shown. The thick and thin lines in the violin plot represent the medians and quartiles of each group, respectively.

**Figure S3.**
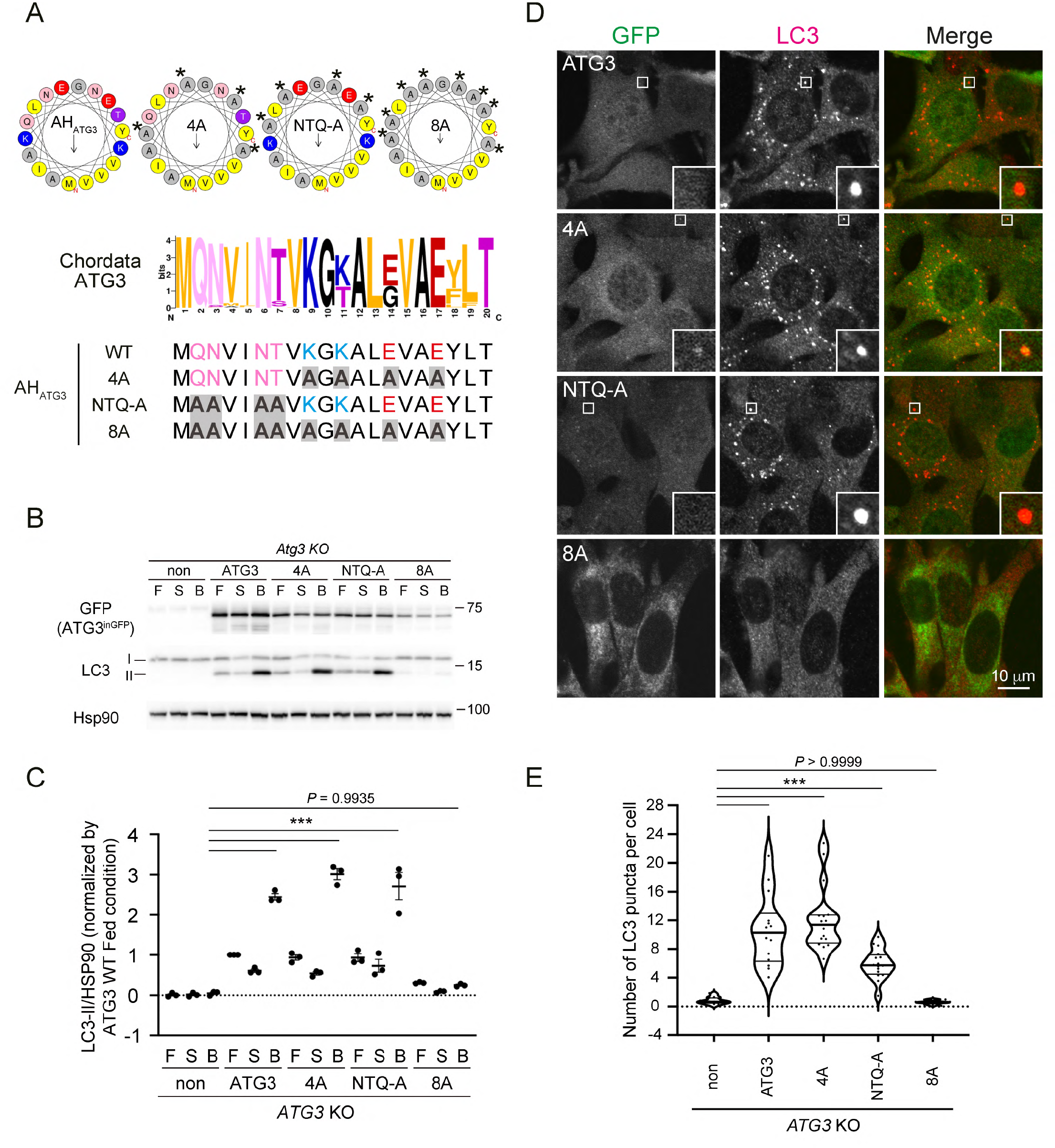
related to Figure 3. Requirement of polar residues in the hydrophilic face of AH_ATG3_. (**A**) (top) Helical wheel representations of AH_ATG3_ mutants containing mutations in its hydrophilic face. Asterisks indicate the position of mutations. (middle) WebLogo to represent amino acid sequence conservation among AHs of Chordata species’ ATG3 proteins. (bottom) Multiple sequence alignment of AH regions of ATG3 WT and mutants. The positions of mutation are marked by gray shadow. (**B**) LC3 flux assay of *ATG3* KO cells expressing ATG3 WT or the indicated ATG3 mutants. The cells were starved for 6h with (B) or without 100 nM Bafilomycin A_1_ (S) or cultured in full media (F). Cell lysates were analyzed by immunoblotting using the indicated antibodies. (**C**) Band intensity quantification of LC3-II shown in Fig. S3B. All data were normalized with those of HSP90. Data represent the mean ± SEM of three independent experiments. (**D**) LC3 puncta formation. The cells were starved for 1 h, fixed and stained with anti-LC3 antibody. The specimens were analyzed by FV3000 confocal microscope. Scale bar, 10 μm. (**E**) Quantification of the number of LC3 puncta. The thick and thin lines in the violin plot represent the medians and quartiles, respectively. The average number of LC3 puncta per cell was counted from randomly selected areas (n≧14). Differences were statistically analyzed by one-way ANOVA and Turkey multiple comparison test. \*\*\**P* < 0.001.

**Figure S4.**
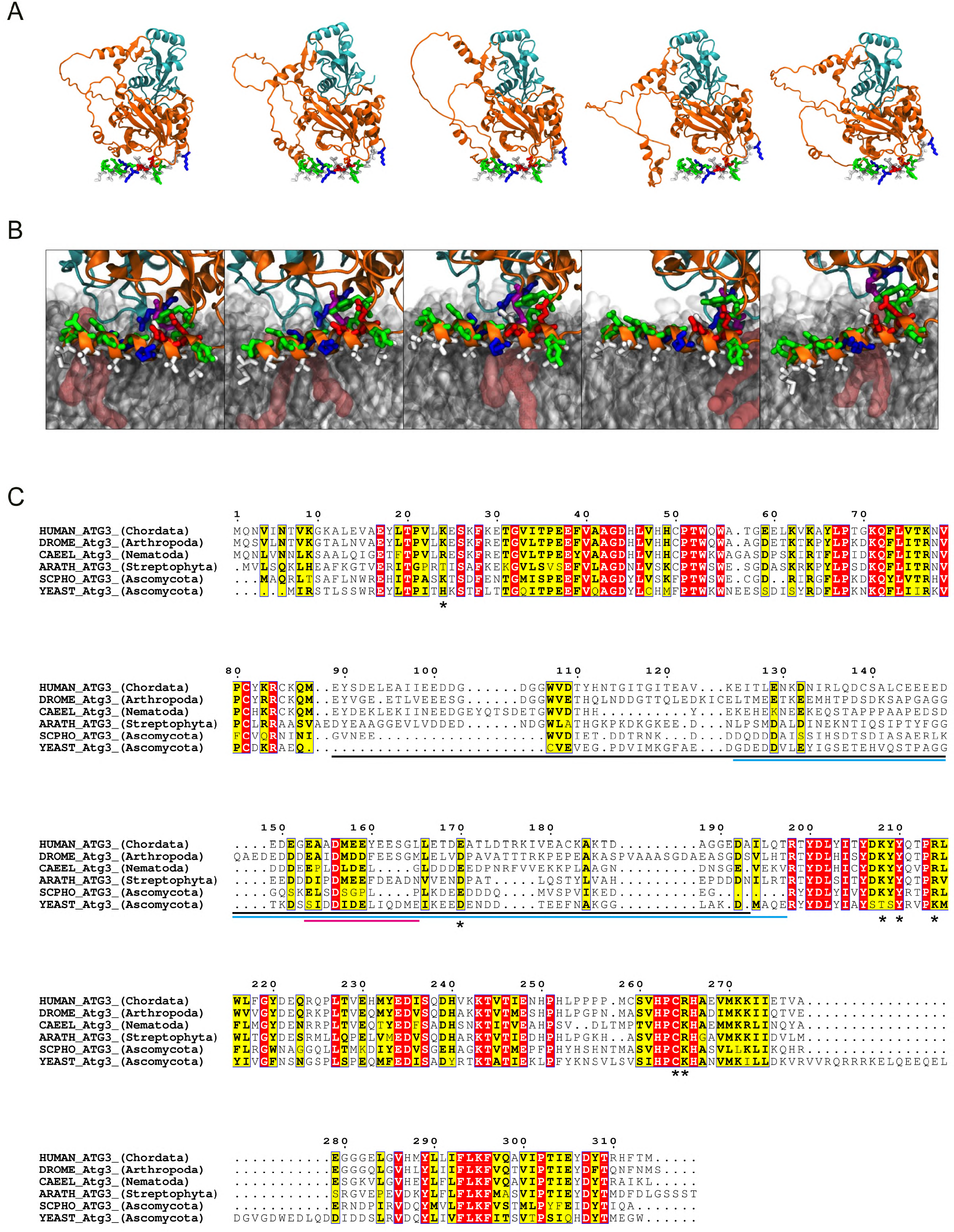
related to Figure 4. Alpha Fold models and AH_ATG3_ dynamics in the bilayer. (A) Five alternative initial models of the ATG3∼LC3 conjugate produced by Alpha Fold. The renders show the complex in a cartoon representation based on its secondary structure, with Atg3 in orange and LC3 in cyan. The atoms of residues 1-24 (AH and linker) are also shown explicitly in a licorice representation. (**B**) Renders of representative snapshots from a trajectory illustrating the dynamics of the ATG3 AH in the bilayer. Rendering and colors as in (**A**). Lipids are showing in transparent grey. A PE lipid forming a contact with LC3 is highlighted in burgundy. (**C**) Sequence alignment of ATG3 proteins using ESPript 3. The identical and similar residues are highlighted by red and yellow boxes, respectively. The flexible region, unfolded region and ATG12-binding region are indicated by black, cyan and magenta lines, respectively. Asterisks show the position of lipid-binding residues *in silico*.

**Figure S5.**
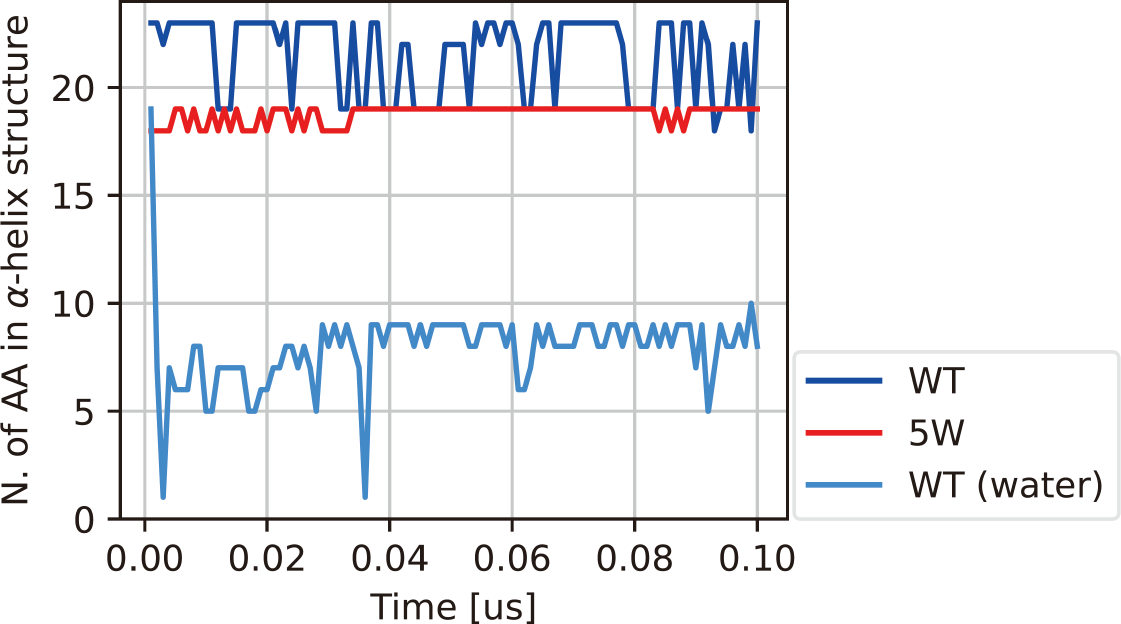
related to Figure 5. Stability of the AH and compactness of the ATG3∼LC3 conjugate Time series of the number of AH amino acids that are helical for WT and 5W in bilayer, and for WT in water.

**Table S1.**
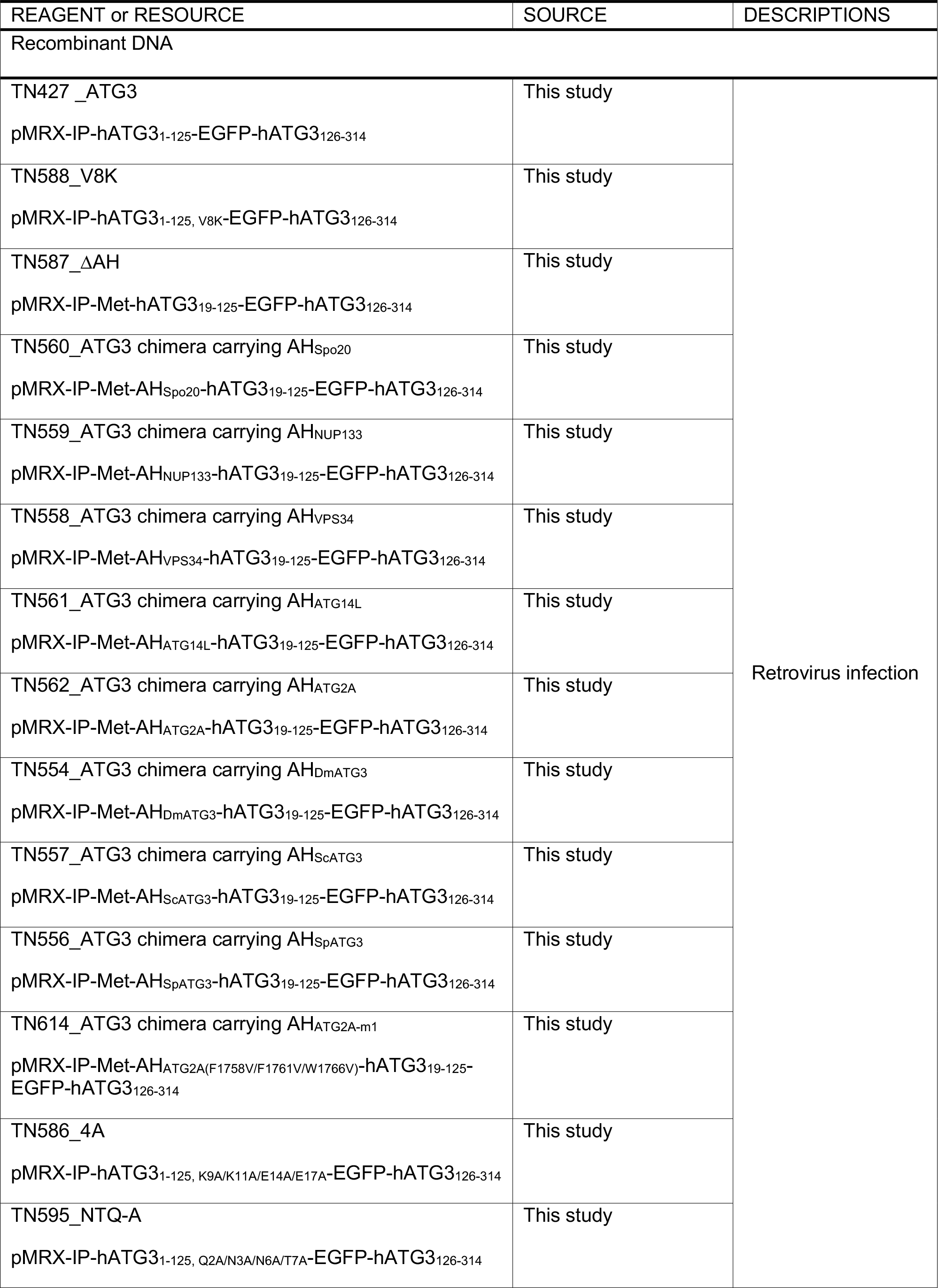
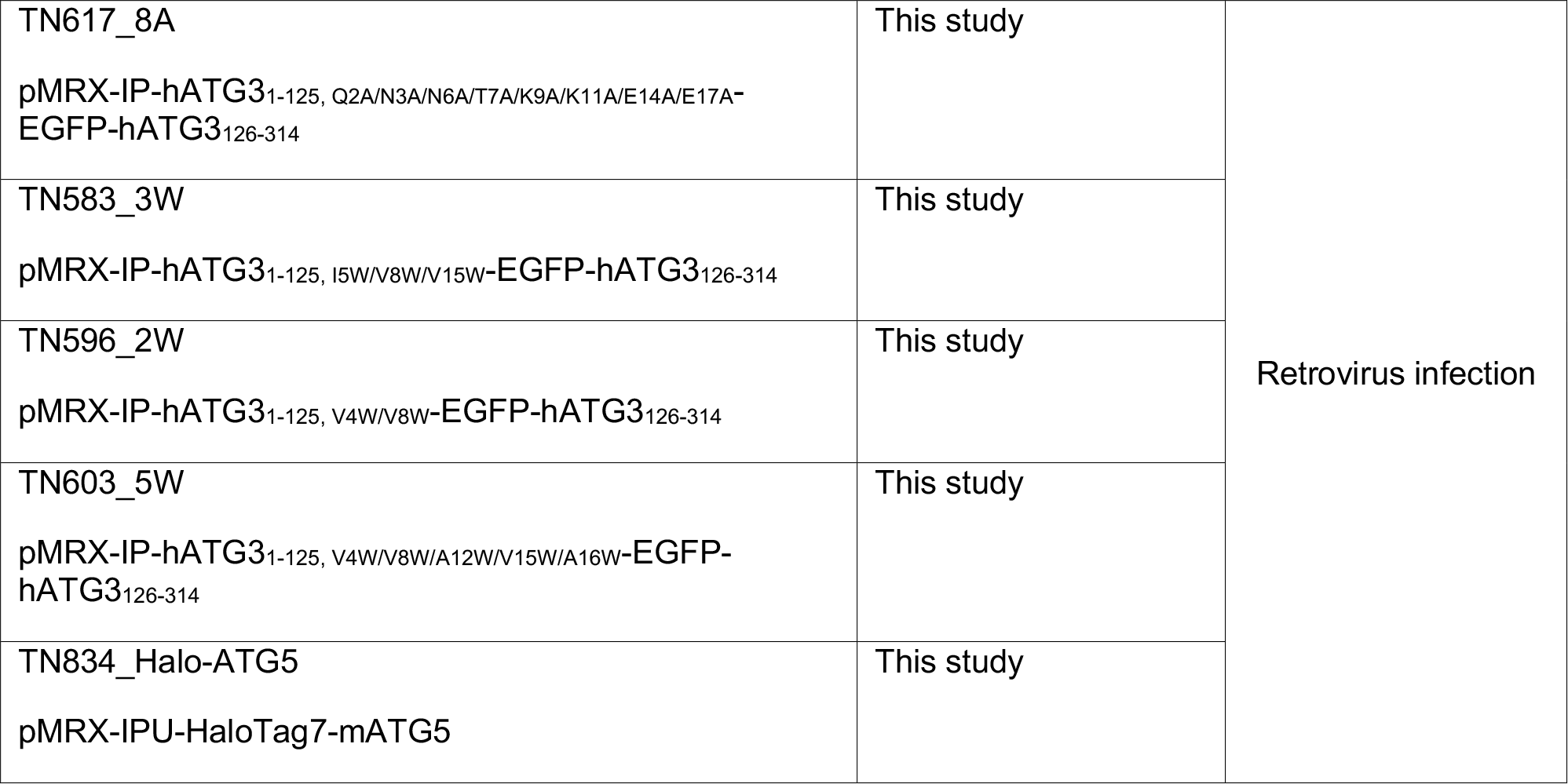
Constructs used in this study

**Movie 1**. Live-cell time-lapse images of ATG3-inGFP and Halo-ATG5 under starved condition.

**Movie 2**. Live-cell time-lapse images of ATG3-inGFP and Halo-ATG5 under starved condition.

**Movie 3**. Live-cell time-lapse images of ATG3ΔAH-inGFP and Halo-ATG5 under starved condition.

**Movie 4**. Live-cell time-lapse images of ATG3(3W)-inGFP and Halo-ATG5 under starved condition.

**Movie 5**. Live-cell time-lapse images of ATG3(5W)-inGFP and Halo-ATG5 under starved condition.

